# Influenza A virus RNA Polymerase targets the chromatin of innate immune response genes

**DOI:** 10.1101/2022.03.30.486363

**Authors:** Jia Yi, Jessica Morel, Mickaël Costallat, Nathalie Lejal, Bernard Delmas, Christian Muchardt, Eric Batsché

## Abstract

**KEY POINTS:**

- FluPol is linked to RNAs in gene body and 3’-downstream regions
- FluPol Chromatin-targets are involved in the cell defense
- Cap-snatching could occur everywhere during transcription elongation

The influenza A virus (IAV) RNA polymerase (FluPol) primes the viral transcription by using capped 5’-ends « snatched » from host nascent RNAs. However, the exact localization of the FluPol on the genome or the timing of its snatching activity remains poorly characterized. Here, we have monitored IAV infection from the perspective of the FluPol interaction with the host chromatin template. Quantification of chromatin-bound RNAs shows a significant perturbation in host transcription that correlates with a relocalization of RNA polymerase II (RNAPII) from the gene bodies to downstream intergenic regions. This extended transcription leading to the production of RNA downstream of genes (DoGs) was previously linked to the NS1-mediated inhibition of the transcriptional termination. However, immunoprecipitation of FluPol-bound RNA in the chromatin fraction revealed that FluPol remains linked to nascent host transcripts during phases of transcriptional elongation and termination, thereby extending the window of opportunity in which cap-snatching may occur. In addition, chromatin-associated FluPol was enriched at transcription termination sites, suggesting that it may participate in the virus-induced termination defects. Finally, we observed that, rather than targeting just highly expressed genes, the FluPol was preferentially recruited to promoters activated by the viral infection and enhancers. Together, these observations suggest the FluPol uses the early immune response to target regulatory elements of defense-related genes at which it interferes with the fate of the transcripts, and possibly also limits RNAPII re-initiation by impairing the termination process.

**Author Summary:** The influenza A virus (IAV) exploits the transcriptional machinery of the infected cells to generate its own RNAs with the viral polymerase (FluPol) hijacking cap structures from host mRNAs. This “cap snatching” requires an interaction between FluPol and the cellular RNA polymerase (RNAPII) accumulating at transcription start sites (TSS) of genes. In parallel, another IAV factor interferes the termination of gene transcription, resulting in frequent transcription downstream of genes and reduced recycling of the RNAPII. Here, employing genome-wide ChIP assays, we show that FluPol selectively targets the TSS of infection-activated genes often associated with cellular defense mechanisms, and thereby contributing to their downregulation. In addition, we show that FluPol is not only recruited to promoters and enhancers, but also within genes, where it interacts with nascent RNAs during transcription and splicing. It is further detected beyond the 3’ end of the genes, suggesting that FluPol exerts a broad influence on host transcription by affecting both transcription initiation through cap-snatching and transcription re-initiation via its involvement in the virus-induced transcription-termination defect at the end of genes.

## Introduction

Influenza A virus (IAV) is an RNA virus whose replication occurs in the nucleus of infected cells (Herz et al. 1981) and its own transcription is dependent on activity of the cell RNA-polymerase II (RNAPII) [1]. This feature, atypical for RNA viruses arise from the activity of the viral RNA polymerase complex (FluPol) formed by the intricate association of the PA, PB1 and PB2 subunits. Specifically, FluPol requires RNA primers originating from the endonucleotidic cleavage of the 5’-end of capped cellular transcripts by the PA subunit to produce viral transcripts [2–4]. This process called ‘‘cap snatching’’ allows FluPol to produce viral translation-competent mRNAs by combining the 10-15 first nucleotides from a capped host RNA with a viral RNA sequence generated by copying the negative sense genomic vRNA (-). Deep-sequencing of the host-encoded portion of these chimeric mRNAs has indicated that about half of these sequences matches 5’-ends of mRNAs [5] or snRNAs in proportion to their transcription level [6].

Several observations suggest that FluPol steals capped RNAs in the early phases of transcription elongation, close to the promoters, in the region where RNAPII is pausing before entering the gene [7]. Firstly, FluPol preferentially interacts with RNAPII phosphorylated at the serine-5 (S5p) within the repeated motifs of the C-terminal domain (CTD) [8–10]. Analysis by mNET-seq of viral RNAs co-immunoprecipitated with the RNAPII has further shown a 1000-fold enrichment in S5p-RNAPII compared to the total pool of RNAPII, suggesting that FluPol generating the vRNA segments requires the association with S5p-RNAPII [11]. This form is enriched at promoters as the S5 phosphorylation event activates the capping reaction, a required maturation step preceding the elongation of nascent RNAs [12–15]. Furthermore, in mass-spectrometry-based approaches, FluPol was found to interact with transcriptional co-regulators enriched at promoters, such as DDX5, coAA/RBM14, PSF/SFPQ or HDAC1/2, and with the cap-promoting elongation factor DSIF (SPT4/5) [16–19]. Likewise, FluPol interacts with CHD1 [20], a chromatin remodeler involved in transcriptional elongation and binding to histone 3 tri-methyled at lysine 4 (H3K4me3), an epigenetic hallmark of active promoters [21].

Finally, an interaction between FluPol and the nuclear RNA exosome complex was reported to favor the recruitment to promoters [22]. Cap-snatching at early phases of transcription also provides an attractive explanation for the coupling of the vRNAs synthesis with the activity of the RNAPII [9], ensuring FluPol to find the capped-RNA primers. Existing observations suggest only minimal or no specificity in the selection of promoters that act as cap-snatching sites and, as a result of a basic rule of probability, it seems to occur more frequently in highly transcribed mRNAs [5,22], U snRNA and snoRNA genes [6,23].

Some observations suggest however that the model of “non-specific, promoter-only targeting” may not entirely reflect the complexity of the interaction between the viral and the cellular transcription machineries. Firstly, the limited depth of the RNA sequencing and the challenge of unambiguously map the very short snatched host primers (10-15 nucleotides) may be a major source of bias that could explain, at least in part, the apparent tropism of FluPol for highly expressed genes. Next, while S5p-RNAPII is enriched at promoters, it is not absent from the body of genes. Likewise, capping enzymes are not only recruited at transcription start sites (TSS), but are also present inside genes at the transcriptional termination sites (TSEs) and in the 3’ flanking downstream regions [24]. Capping factors could, therefore, produce capped RNAs not only at TSSs but potentially at any location where RNAPII is transcriptionally active, as evidenced by the capping of short non-coding pervasive transcripts detected within genes and at both 5’ and 3’ ends [25,26]. Capped RNAs are also produced by enhancers, which are actually more abundant on the genome than promoters [27–29]. Thus, if the sole purpose of FluPol recruitment to chromatin is cap-snatching, then a wider distribution than just promoters would be expected. Finally, the cap-snatching activity of FluPol is considered a component of the viral strategy to evade host cell defense, as cap loss leads to mRNA destabilization and altered protein synthesis. Yet, IAV also interferes globally with host transcription by causing a termination defect. By inhibiting the RNAPII termination complexes through the action of its NS1 factor [11,30], it disrupts the cohesin/CTCF-anchored chromatin loops [31], resulting in the opening of host topologically associating domains (TADs). This causes long run-through events of RNAPII in intergenic regions, technically capturing the polymerase on chromatin and preventing re-initiation at promoters. This extended transcription leads to the production of RNA downstream of genes (DoGs), a phenomenon also observed in cell undergoing stress [32,33]. Like the cap-snatching, this strategy also occurs on the chromatin and involves the transcription machinery, and it is tempting to speculate that these seemingly separate strategies could be two facets of a same mechanism.

We therefore deemed it necessary to reinvestigate the crosstalk between the host- and viral-transcription machineries. To this end, we have monitored the evolution of chromatin-associated transcripts upon IAV infection, while also following the recruitment of FluPol to the chromatin. For the latter, we performed a series of chromatin immunoprecipitations (ChIPs) using different antibodies recognizing the FluPol subunits PA, PB1 and PB2. We also transduced a FluPol PA subunit labeled with a highly efficient V5 epitope to perform ChIP assays and analysis of chromatin-bound RNAs.

This unprecedented approach surprisingly revealed that FluPol remains linked to the nascent transcripts (probably through the RNAPII association) during phases of transcriptional elongation and termination. Additionally, we found that genes recruiting FluPol to their TSS, TES, gene body, or enhancers were more frequently involved in cellular defense than expected by chance. Thus, our findings collectively indicate that FluPol associates with virus-activated genes starting from their TSS and extending all the way to their downstream 3’ end region. We therefore propose an impact of FluPol on host cell transcription not only at the initiation phase, but also during elongation and termination, thereby maximizing its dampening effect on host cell defense mechanisms.

## Results

### High-throughput sequencing of chromatin-associated RNAs in IAV-infected cells revealed extensive transcriptional perturbations

To examine the impact of influenza virus infection on host transcription without the confounding effects of RNA maturation, we sequenced chromatin bound RNA in human lung epithelial A549 cells infected at a high MOI (MOI = 5) with an H1N1 influenza virus (A/WSN/33). At 6 hours post infection (hpi), we observed that up to 40% of the 75% most expressed genes were differentially regulated, as evaluated by the DESeq2 package. (**Fig 1A and S1A Fig**). Thus, the number of dysregulated genes was even greater than what has been previously reported with total RNA extracts at early times of infection [11,31,30,34–36].

**Figure 1:**
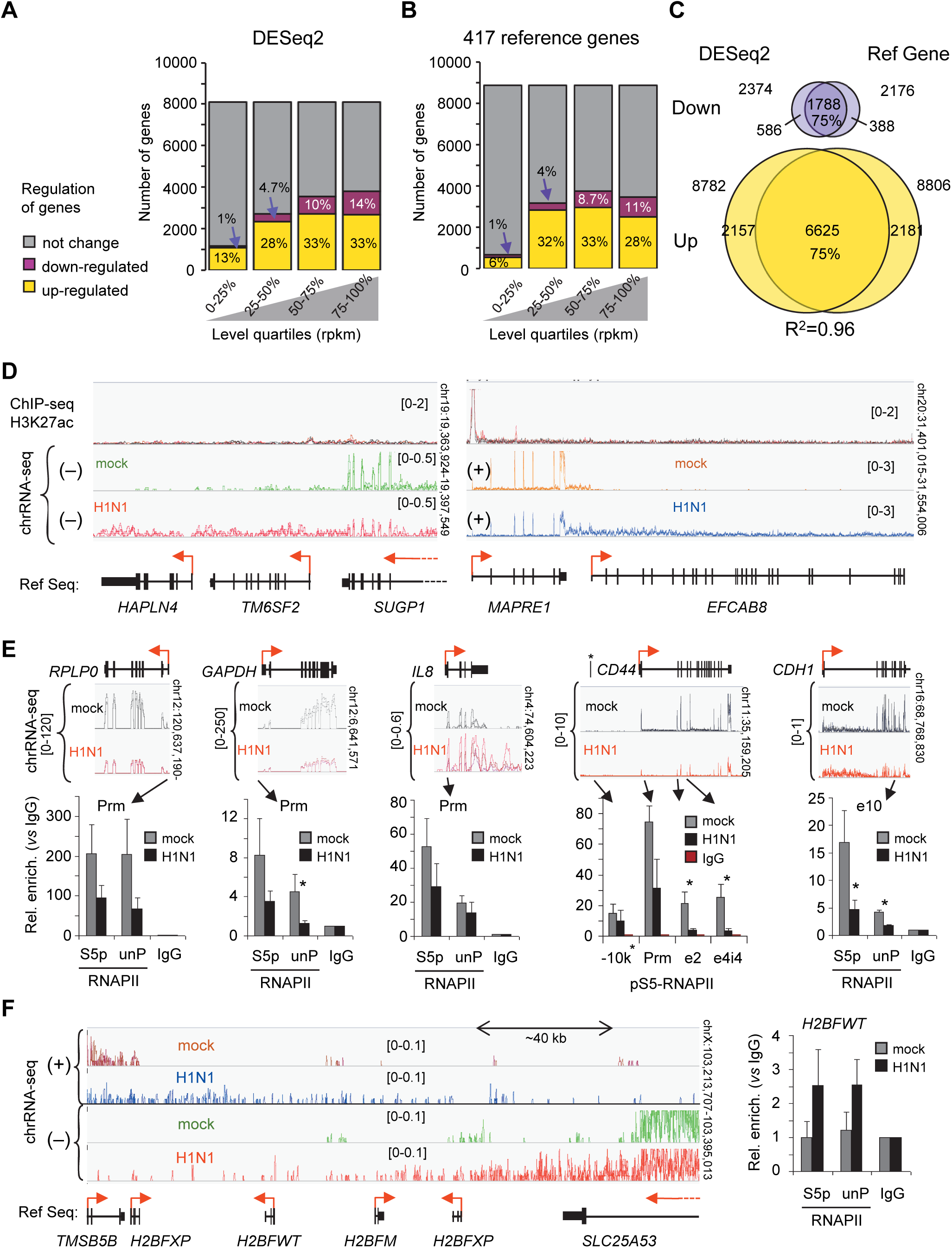
Chromatin-associated RNAs illustrate cell transcriptional changes induced by 6 h of IAV H1N1 infection in A549 cells. **A**) Differential RNA levels between H1N1-infected cells and uninfected cells (mock) using DESeq2 package. Genes were considered regulated if the fold change was >1.5 and the adjusted p-value was <0.05. Genes were ranked in quartiles as indicated. **B**) Differential RNA levels using normalization based on the average variation of 417 reference genes expressing higher levels (> 300 rpkm) of transcripts and with long half-lives (> 15 h) [37] from a paired t-test (two-tailed). Genes were ranked in quartiles from the mean of the two conditions. **C**) Venn diagram for up- and down-regulated genes evaluated by the two methods. R2 is the Pearson’s correlation coefficient comparing the two methods. **D**) Examples illustrating the variation in read distribution in H1N1 infected cells from IGV visualization of indicated loci. The RNA-seq independent triplicates have been overlaid in the same color as indicated for each strand orientation. Top track shows H3K27ac enrichment on chromatin from infected or mock cells. Track scale ranges are indicated in brackets. The bottom track (Ref Seq) shows the position of the genes and their orientation as indicated by the red arrows. **E, F**) ChIP assays using antibodies against S5 phosphorylated (S5p) or unphosphorylated (unP) RNAPII from infected and uninfected cells. Immunoprecipitated DNA was quantified by qPCR using primers targeting the indicated regions. Enrichment is expressed relative to the signal obtained for ChIP with negative IgG control. Values are mean (± deviation) of at least three independent experiments. Statistical significance of differential levels between infected and non-infected cells was evaluated by Student’s t-test (two-tailed), with P < 0.05(*).

As previously reported, the strong perturbation of the cellular transcriptome induced by the IAV infection results in a substantial redistribution of RNA-seq reads both in-between genes and between genes and intergenic regions [11,30]. Such a heterogeneous transcript distributions is known to affect the performance of statistical normalization [38–40]. Therefore, we used an alternative normalization method based on a set of reference genes. We reasoned that pools of transcripts with long half-time would remain largely unchanged after only 6 hours of infection. Thus, considering that the 20% most expressed transcripts (> 300 rpkm) with long half-lives (> 15 h) [37], we identified that approximately 40% of the 75% most expressed genes were differentially regulated (**Fig 1B, S1B Fig**), but with only a 75% overlap with the genes identified by DESeq2 pipeline (**Fig 1C, and S1 Table**). A gene ontology analysis on genes up-regulated at least 4-fold (p-value<0.05) upon infection confirmed the anticipated enrichment in pathways involving type-I interferon, cytokines, and OAS ribonuclease family members (**S1C Fig, S2 Table**).

Earlier studies have shown that IAV-dependent transcriptional termination defects result in the production of ‘‘Downstream-of-Gene’’ transcripts (DoGs) at many active genes [11,30,31]. Sequencing chromatin-bound RNAs proved very efficient at visualizing DoGs and RNAPII terminal read-through was detected at essentially all genes relying on the canonical polyadenylation complex, including several genes previously reported as unaffected (*LY6E*, *APOL1*, *DEFB1* and *IFI6*) [30] (**S1D Fig**). Consistent with a role of the polyadenylation complex, and in agreement with earlier observations [41,11], replication-dependent histone genes did not produce DoGs (**S1E Fig**). This very general termination defect may largely contribute to the bias we corrected above, as we observed a general decrease in apparent exon coverage associated with an increase in the abundance of reads beyond the site of polyadenylation (**S1F Fig** and **Fig 1D**).

To gain further understanding of the mechanisms perturbing transcription, we carried out RNAPII ChIP on infected cells. This revealed that the decreased exon coverage in the RNA-seq data was concomitant with a decreased RNAPII accumulation at the promoters of highly expressed genes either unaffected (*GAPDH*, *RPLP0*) or induced (*IL8*) upon infection, and on the transcribed region of genes moderately expressed (*CD44*, *CDH1*) (**Fig 1E**). Inversely, we noted an increased accumulation of RNAPII at loci corresponding to genes not expressed in A549 cells even after IAV infection, such as *H2BFWT*, possibly as a consequence of a read-through from neighboring genes (**Fig 1F**). Re-analysis of publicly available ChIP-seq data from primary human monocyte-derived macrophages (MDMs) infected with IAV and harvested at 6 hpi [31] revealed a very similar decrease in RNAPII recruitment at expressed genes associated with an increased accumulation on intergenic regions and at genes not expressed in these cells (**S1G Fig).**

### Quantification of chromatin-bound RNAs confirms that the IAV infection induced defects of transcriptional termination

We next reexamined the DoGs in the light of their improved detection offered by the sequencing of chromatin-associated RNA. Visual examination in a genome browser showed that numerous genes produced DoGs in the absence of IAV infection, illustrating the intrinsic imperfections of the termination machinery (see examples **Fig 1D, S1D S1F Figs)**. As production of DoGs has been proposed to participate in the global reduction in host cell transcription, we examined the correspondence between gene expression and production of DoGs. To assess alterations in DoG production in infected cells, we quantified the density of oriented reads in 10-kb windows within a 40 kb region downstream of TESs (**Fig 2A)**. This quantification showed a robust overall upregulation of DoGs even at sites distal from the TESs within the 40kb region under scrutiny (**Fig 2B)**. In total, 44% of the asserted downstream regions displayed transcription upregulated 1.5-fold or more, while down-regulation was observed at less than 1% of the regions (**Fig 2C)**. Upregulated DoGs originated from up-regulated genes in approximately 30% of the cases, suggesting that production of DoGs does not *per se* prevent gene activation, at least at early times of infection. Yet, the majority of the upregulated DoGs (∼55%) were located downstream of genes unaffected by IAV infection (grey proportion of up in **Fig 2D**), strongly arguing in favor of the previously suggested impact on termination. Finally, we surprisingly noted that approximately 10% of the upregulated DoGs, as well as 70% of the non-significantly modified DoGs, were associated with silent genes. Through visual verification, we were able to provide an explanation for this phenomenon, as we observed that these particular cases involved genes that were entirely traversed by DoGs originating from neighboring genes.

**Figure 2:**
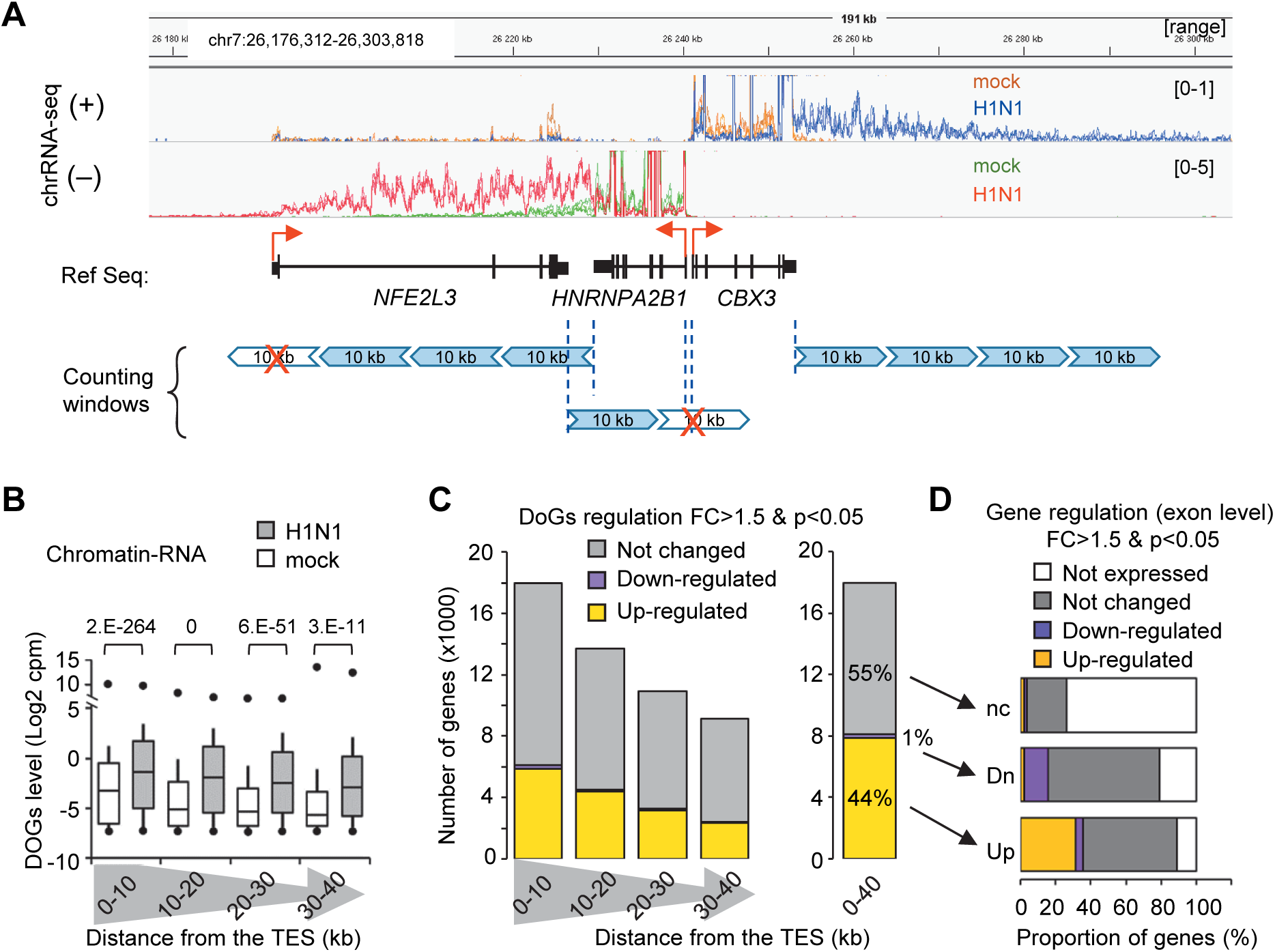
Chromatin RNAs were enriched in DoGs within the 3’-end region of the genes after IAV infection. **A)** Strategy to quantify the 3’-end transcripts downstream of the termination sites (TES). Oriented reads were counted in 10 kb windows downstream to the TES extending until 40 kb. The downstream-10 kb windows which overlapped with the transcription start site of another Ensembl gene in both orientation were not taken in account (red crosses). **B)** For each selected gene, the indicated 10-kb windows from the TES were quantified for their DoGs levels evaluated in Log2(cpm). Data are presented in box plots where the bottom line is the 1st centile, the box corresponds to the 2nd and 3rd quartiles, the top line is the 9th centile, and the middle line in the box is the median. Small circles indicate the minimum and maximum values. P-values indicated at the top evaluate the statistical significance of the difference in levels between infected and uninfected cells as calculated by Student’s t-test (two-tailed). **C)** Number of genes for which a 10-kb window downstream of the 3’-end showed a differential transcription level. These windows were considered modified for a p-value<0.05 (paired t-test, two-tailed) and a fold change >1.5. The right panel shows the same quantification for all pooled windows. Transcription downstream of TES covering the 40 kb was considered upregulated if the majority of the 10-kb windows were upregulated (yellow), and vice versa for downregulation (magenta). The 40 kb region was considered unchanged (gray) if the 10 kb windows themselves were found to be unchanged, or if there was no significant difference in the level of DoGs on the four 10 kb windows, as evaluated by a paired t-test (two-tailed, p<0.05). **D)** For each regulated category (not changed nc, down-regulated Dn, up-regulated Up) of the 3’-end downstream region, the regulation of the corresponding genes (as evaluated in **Fig 1B**) is indicated in proportion.

### Genome wide Analysis revealed recruitment of FluPol at innate immune genes

We next explored the genome-wide recruitment of FluPol by chromatin immunoprecipitation-sequencing (ChIP-seq) using antibodies directed against the PA, PB1, and PB2 subunits of the viral polymerase. These antibodies, when tested on extracts from infected cells, all co-immunoprecipitated phosphorylated RNAPII (**S2A Fig**), consistent with earlier observations documenting an interaction of the FluPol with the pS5-RNAPII) [8–10].

To avoid interference from neighboring genes, we focused our analysis of the ChIP-seq data at genes located at least 5 kb away from other genes. Specific examination of the ChIP signals on 500 bp-windows centered on promoter TSSs, showed a significantly increased signal upon viral infection, which was more robust at expressed genes (as selected on the read-count from the RNA-seq) than at unexpressed genes **(Fig 3A)**. The specificity of the assay was further supported by the increased signal observed at active promoters harboring H3K27ac marks, that was more important than at TSS of unexpressed genes (compare middle graph to the right one, **Fig 3A**). The differential enrichment at TSS induced by the infection for each subunit was significantly greater on expressed than on unexpressed genes (above the red line, **Fig 3B**).

**Figure 3:**
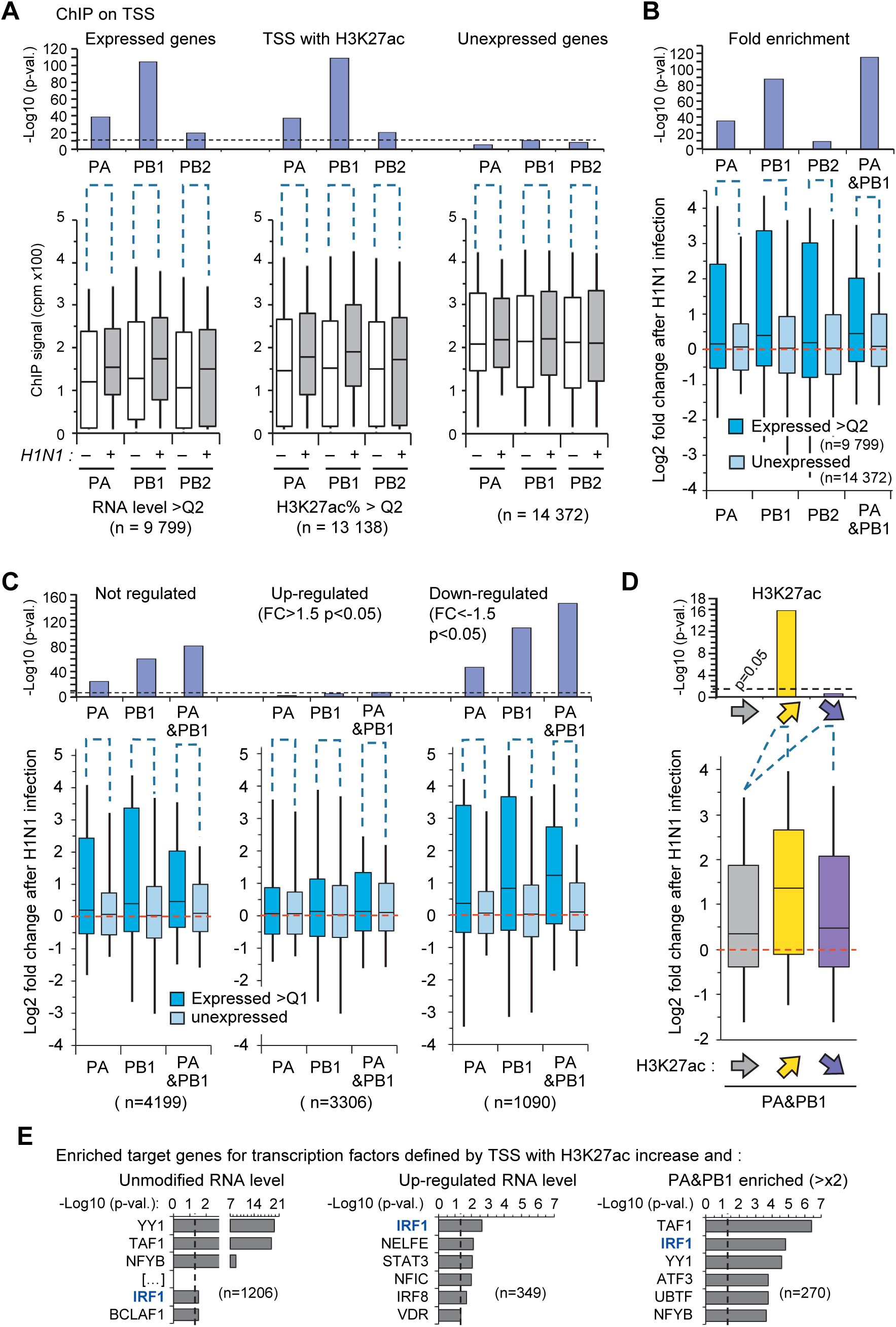
Enrichment of FluPol subunits on the chromatin. Chromatin from uninfected (-) or infected (+) A549 cells was immunoprecipitated with the indicated subunits of FluPol and analyzed by deep sequencing. For each 500 bp window centered on annotated transcription start sites (TSS), DNA enrichment was the mean of replicates expressed in cpm (multiplied by 100 for ease of presentation). Only TSSs separated from other TSSs by 5 kb were considered to avoid potential bias from close neighbors or overlapping genes. **A**) Bottom box plots illustrate the distribution of TSS from genes expressed above the median (> Q2), TSS with H3K27ac signal greater than the median of H3K27ac signals, or TSS from non-expressed genes. The bottom and top lines of the box plots indicate the 1st and 9th centiles, respectively. The upper graphs show the p-values evaluated on log2 normalized cpm by a paired t-test (two-tailed) assessing the enrichment induced by H1N1 infection on each TSS in the indicated categories. The dotted line indicates the p-value threshold based on the unexpressed genes. **B**) The fold change of enrichment for the indicated TSS categories are the averages of the TSS ratio of infected versus non-infected cells for the signal of FluPol subunits. Statistical analysis was performed on log2 fold change using Student t-test (two-tailed) to compare enrichment on TSS of unexpressed genes (light blue) and genes expressed above the median (blue). The dashed red line indicates no change. The p-values are given above the box plots. In each case, the number (n) of TSS tested is indicated. PA&PB1 analyses were performed by combining their ChIP-seq signals..**C**) Fold change of enrichment for TSS of genes in indicated categories. As described in B) Enrichment of the indicated FluPol subunits on the TSS of the 75% of the most highly expressed genes that are regulated by more than 50% (p<0.05) or are not regulated. Enrichments were compared to the signal of FluPol subunits on TSSs of unexpressed genes using a Student t test (two-tailed) to calculate p-values. **D**) Fold change of enrichment of combined PA&PB1 signals on TSS with increase (yellow, n=804) or decrease (violet, n=716) of H3K27ac content. H3K27ac levels were normalised to H3 levels and were considered regulated when the 75% most enriched TSS/H3K27ac changed by more or less than 20% with p<0.05. These enrichments were compared to those for TSS with unmodified (grey) H3K27ac with the same H3K27ac levels to calculate p-values by unpaired student t-test (two-tailed). **E**) Enrichment of genes with consensus of indicated transcription factors in ENCODE and ChEA, which are ChIP-seq enrichment analysis tools built from gene set libraries generated from published ChIP-seq data extracted from multiple sources. Genes were selected based on their TSS containing H3K27ac up-regulated by more than 20% (p<0.05) and either their RNA level change as indicated or PA&PB1 recruitment in infected cells (2-fold more than mock cells, p<0.05). P-values of these gene enrichments were evaluated using Enrichr and the threshold of p<0.05 is indicated by the dotted line.

We further noted that the combination of PA and PB1 signals was more efficient at detecting expressed genes than each signal taken individually, designating this combination as a good approach for detection of FluPol (right bars in **Fig 3B**). In contrast, the anti-PB2 antibody behaved relatively poorly in discriminating the specific signal induced by the infection on expressed genes versus unexpressed genes (**Figs 3A, 3B**). Finally, in introns, we observed no difference in the recruitment of FluPol subunits between expressed and silent genes (**S2B Fig**). Together, these observations were consistent with a recruitment of the FluPol subunits to sites of transcriptional initiation.

To investigate whether FluPol recruitment was also governed by criteria other than transcriptional activity, we further carried out a Gene Ontology analysis on the genes recruiting PA and PB1 by at least 4-folds on TSS. This revealed a significant enrichment in genes associated with cell signaling, including kinase and membrane trafficking factors. More importantly FluPol was found on the TSS of several genes of the innate immune system such as *ILF2*, *MTDH* enabling NFkB activity and factors with double-strand RNA-binding activity [42] (**S2C Fig**), *SIDT2* mediating the long dsRNA entry in lysosomes for degradation [43], and other dsRNA detectors such as *OAS1/2/L*, *DDX58/RIG-I*, *DDX60*, *DHX36* and *DHX30* (**S2G Fig** and **S3A Table** for a full list). The outcome of the pathway analysis of FluPol target genes (**S2F Fig, S3A Table**) diverged at least in part from that reached when examining highly expressed genes, which were identified based on either an elevated H3K27ac levels at their promoter (upper centile, **S2D Fig**), or a high RNA-seq signal (upper quartile, **S2E Fig)**. The pathways defined as RNA binding (GO:0003723) and double-stranded RNA binding (GO:0003725) were commonly shared between lists of FluPol-bound and highly expressed genes (**S2F Fig, S3A Table**).

However, comparison of the GO term enrichments revealed 17 specific pathways significantly enriched for FluPol genes and not significantly enriched for genes with the highest H3K27ac or RNA levels. (**S2F Fig)**. For instance, FluPol preferentially targeted TSSs of genes related to tRNA modifications (GO:0140101, such as *TARS2* and *DUS1L* for example, **Fig 3G**). This suggested that FluPol enrichment at TSS can only in part be accounted for by promoter activity, while gene function also seems to play a role particularly at genes involved in cell defense and protein synthesis.

To investigate the potential impact of FluPol recruitment on gene expression, we compared the TSSs enriched in viral polymerase with genes categorized by their differential expression upon IAV infection. Surprisingly, up-regulated genes were the less significantly enriched in FluPol, being out performed by downregulated or unaffected genes (**Fig 3C**). This observation supported the hypothesis that FluPol’s Cap-snatching function reduces gene expression since up-regulated genes had low FluPol recruitment while down-regulated genes had high recruitment. Hence, the set of genes that did not show significant changes might contain those that are typically upregulated upon activation of cellular defense mechanisms but effectively suppressed by FluPol. Accordingly, the FluPol recruitment was higher on TSS with increased H3K27ac levels compared to the recruitment on promoters with decreased H3K27ac levels (**Fig 3D**). This suggested that FluPol may exhibit a better affinity for activated promoters (with H3K27ac increase) and this may contribute to the decrease in gene expression, especially at genes involved in the innate immune response. It is noteworthy that among the FluPol targeted promoters showing an increase in H3K27ac, only the IRF1 target genes were enriched in upregulated genes and in unchanged genes. (**Fig 3E, S3B Table**). This shows that although the promoter of IRF1 and its target genes are induced (with IRF1 itself exhibiting a 2.3-fold upregulation, as shown in Supplementary Table 1A), the presence of FluPol at the promoters of these IRF1 target genes inhibits their mRNA production.

### FluPol is recruited to enhancers in regions close to immune response genes

Like promoters, enhancers are sites of RNAPII transcription and produce capped RNAs (eRNAs). We therefore investigated whether these regulatory elements would also be sites of FluPol recruitment. To this end, we identified a series of isolated enhancers active in infected cells, based on the presence of the H3K27ac histone mark characteristic of enhancers, and the absence of H3K4me3, enriched at promoters (**Fig 4A** and bioinformatics section in method). FluPol recruitment at these enhancers was then estimated by quantifying the ChIP-seq signal for each subunit in bins of 1 kb around the H3K27ac peak (**Fig 4A**, blue boxes). This approach revealed that, upon infection, all FluPol subunits displayed a significantly increased accumulation within the first 1kb from the center of the enhancer, rapidly decreasing as a function of the distance to the center (**Fig 4B –** in the shown graphics, left and right windows are considered together). As observed for promoters, the anti-PB2 antibody seemed the less efficiently at detecting enhancers. We noted also that the recruitment of FluPol subunits to enhancers could be positively correlated with the extent of their H3K27 acetylation (**Fig 4C**). This suggests the FluPol enrichment on enhancers is dependent on the activity of enhancers.

**Figure 4:**
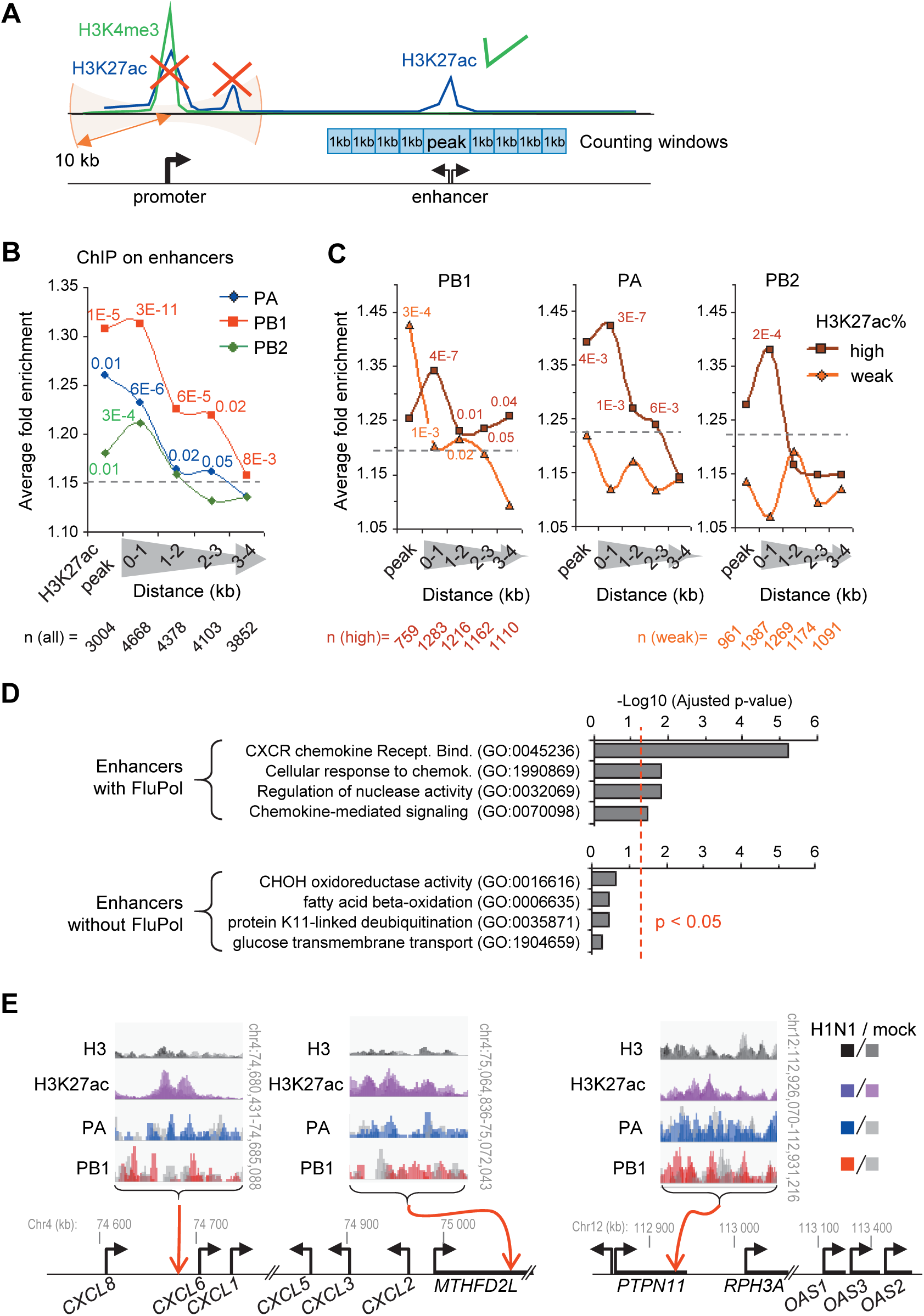
FluPol subunits on chromatin are enriched at the enhancer sites. **A**) Strategy to quantify FluPol enrichment around H3K27ac peaks defining enhancers. Reads were counted in the H3K27ac window and in four 1 kb windows downstream and upstream of the boundaries of the H3K27ac peaks. For each counting window, the fold change enrichment between infected cells vs. mock cells of the indicated FluPol subunits was calculated. The H3K27ac peaks overlapping the 10kb region around the H3H4me3 peaks were not considered (red crosses). **B**) Peaks of H3K27ac detected in infected cells and not overlapping with H3K4me3 peaks ± 10kb defined the first counting windows (named “peak”). Upstream and downstream windows were pooled for each indicated distance. The average of the enrichment ratio between infected cells vs mock cells of the indicated FluPol subunits were shown. **C**) The same calculation was performed for the enhancers with high or low percentage of H3K27ac (versus H3). The enhancer peaks were separated into quartiles of H3K27ac% and the relative enrichment of FluPol subunits is shown for the 4th quartile (high level in brown) and for the 2nd quartile (low level in orange). **B** and **C**) Differences between infected and mock cells were evaluated by paired t-test and significant p-values are indicated. The number of loci considered is indicated above. The height of the dashed lines indicates the fold enrichment limit at which differences are significant. **D**) Analysis of gene regulatory pathways in 1 Mbp regions centered on H3K27ac peaks with enrichment for PA and PB1 subunits (top graph) or without enrichment for PA and PB1 (bottom graph). 603 expressed genes were found around the 97 selected enhancers for which the individual PA and PB1 ChIP signal was significantly increased by at least 50% either on the peaks or on the neighboring windows (0-1 kb) (**S4C Table**). 3208 expressed genes were found around the 1391 selected enhancers to have a percentage of H3K27ac above the median level of all peaks and for the combined enrichment of PA and PB1 to be less than 10% (on peaks and on the 0-1kb windows) (**S4B Table**). Expressed genes from the first 5 genes 500 kbp upstream and from the first 5 genes 500 kbp downstream of the peaks were used for pathway analysis with Enrichr. The adjusted p-value of the significant GO pathways is shown for the FluPol-enriched enhancers, and the first five ranked GO terms are shown for the enhancers without FluPol. The threshold of p<0.05 is indicated by the dotted red line. **E**) Localization of enhancers enriched by FluPol. Examples of enhancers with their indicated neighboring genes to illustrate the pathways in **D**). The ChIP-seq independent replicates performed with antibodies against the indicated proteins in infected or mock cells are overlaid in the indicated color. The black arrows indicate the position of the TSS and the orientation of the gene. Red arrows indicate the localization of the enhancer regions where FluPol enrichment was detected.

We then examined whether enhancers recruiting FluPol were linked to genes with particular functions. To this end, we performed genomic region enrichment analysis, listing genes located in the neighborhood of enhancers recruiting FluPol subunits. When considering enhancers displaying PA and PB1 combined signal at least 20% above background inside the region framed by the 2 first 1kb-windows (a total of 500 enhancers), we identified 1966 expressed genes within a 1 Mbp regions centered on H3K27ac peaks. Ontology analysis on these genes revealed an enrichment in several immune system signaling KEGG pathways found, including the IL-17 signaling pathway (Adjusted p-value = 9E-3) (**S4A Table**). These regulatory pathways were not observed when examining genes located near active enhancers that did not exhibit enrichment of FluPol (**S4B Table**). With a more stringent selection, retaining only enhancers displaying a 50% or more increase in PA and PB1 signals in between the 1kb-windows (a total of 97 enhancers), we identified 603 expressed genes, associated with chemokine-mediated signaling pathways and defense genes with nuclease activity (**Fig 4D** and **S4C Table**). Visual examination of the data also identified PA and PB1–enriched enhancers in the neighborhood of CXCL and OAS genes (**Fig 4E**). These data suggest that the FluPol recruitment on enhancers targets specific regions that could contribute to the immune response by limiting their transcriptional action.

### FluPol is associated with intragenic chromatin-bound RNAs

As an alternative approach to study FluPol recruitment to the host genome, we next implemented an RNA-ChIP approach exploring the range of chromatin-bound RNA species associating with the viral polymerase. As available antibodies were of insufficient quality for this more demanding approach, we generated a V5-tagged version of the PA subunit of FluPol. The resulting recombinant H1N1-derived IAV retained its ability to infect the A549 cells. Immunoprecipitation with an anti-V5 antibody under conditions used for RNA-ChIP (see Material &Methods) on cells infected with the V5-tagged virus, resulted in efficient immunoprecipitation of the tagged-PA (**Fig 5A** and **S3 Fig**).

**Figure 5:**
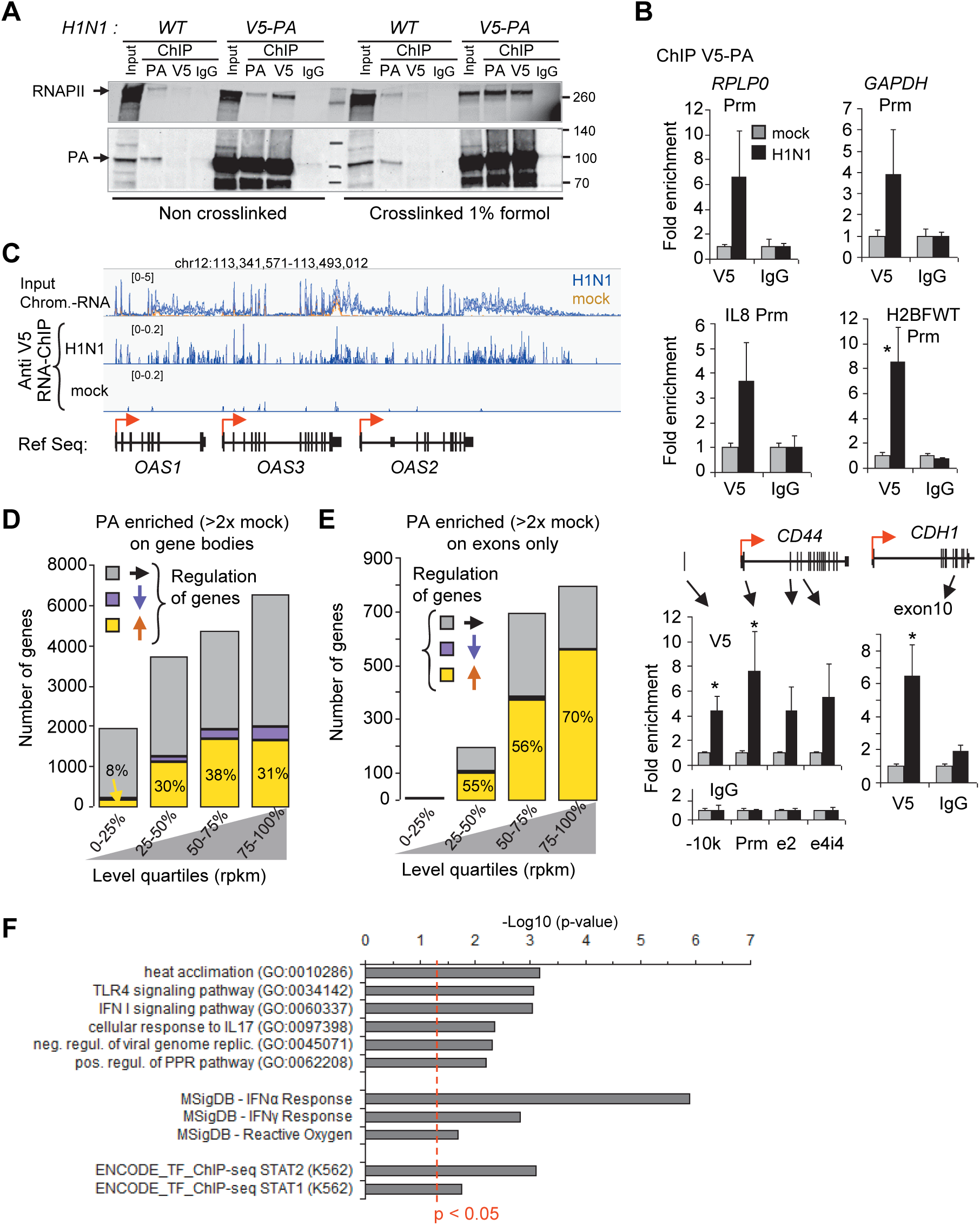
FluPol subunits on the chromatin are associated with nascent RNAs from the host cell. **A**) A549 cells infected with WT IAV or V5-PA engineered IAV were cross-linked by 1% formaldehyde or not. Chromatin fractions were subjected to immunoprecipitation with PA, V5 antibodies or non-immune IgG. Eluates and 10% of extracts were resolved by Western blot. The lower part of the membranes was detected with PA antibodies and the upper part with phospho-RNAPII antibodies.. **B**) ChIP assays using anti-V5 and IgG antibodies from infected and uninfected cells. Immunoprecipitated DNA was quantified by qPCR using primers targeting the indicated regions. Enrichment is expressed relative to the signal obtained for ChIP in uninfected cells. Values are the mean (±dev.) of three independent experiments. Statistical significance of differential levels was evaluated by Student’s t-test (two-tailed), with P < 0.05(*). **C**) Example of PA-enriched RNAs bound to chromatin with IGV. The RNA-seq independent triplicates have been overlaid in the same color as indicated for the (+) strand. The scale range of the tracks is indicated in brackets. The bottom track (Ref Seq) indicates the position of the genes and their orientation by the red arrows. **D**) RNAs from ChIP of V5-PA in V5-PA H1N1 infected cells or in mock cells were analyzed by deep sequencing. PA-enriched RNAs spanning gene bodies were evaluated against IgG and were considered significant if the mean change subtracted from the triplicate deviation was greater than two times that of IgG. The number of enriched genes that were upregulated (yellow), downregulated (magenta) or unchanged (gray) was plotted as a function of gene expression as determined in **Fig 1B** and ranked in quartiles. **E**) The same analysis as in **D**) was conducted for RNAs covering the exons only. **F**) Pathway analysis of 1696 genes on which PA was 4-fold enriched on exonic RNAs between infected and mock cells. P-values of Gene Ontology (GO) biological function 2021, Molecular Signatures Database 2020 (MSigDB), and ENCODE ChIP-seq 2015 were evaluated by Enrichr, and the threshold of p<0.05 is indicated by the dotted red line.

In control ChIP-PCR experiments, the recombinant FluPol containing V5-PA was efficiently detected at the DNA of promoters identified above as recruiting the native version of the viral polymerase, including genes either genuinely expressed at moderate to strong levels (RPLP0, GAPDH, CD44), or induced by the infection (IL8) (**Fig 5B**). Yet, the more efficient immunoprecipitation assays reached with the anti-V5 antibody revealed that V5-PA was also present within the coding region genes (CD44 and CDH1) (**Fig 5B**) or within a read-through region by run-away RNAPII having initiated at promoters of upstream genes (H2BFWT) (**Figs 5B, 1F**).

Implementing RNA ChIP-seq with V5-tagged virus and anti-V5 antibody revealed that chromatin-RNAs associating with V5-PA extensively distributed to the coding region of a large set of expressed genes (a total of 15299 genes, corresponding to 55% of the examined genes; see example of the OAS genes **Fig 5C**).

Quantification further documented that the immunoprecipitated chromatin-bound RNAs were preferentially originating from the most expressed genes, although genes expressed at low levels were also contributing (compare quartiles in **Fig 5D**).

We also identified a smaller set of genes (1809 genes) at which V5-PA-bound RNAs were enriched at exonic sequences (**Fig 5E**). This was indicative of an association of PA with pre-mRNAs having undergone full or partial co-transcriptional splicing, and we will refer to these species as “spliced chromatin-RNAs”. Interestingly, this set of genes was enriched in genes up-regulated by the infection (compare yellow segments in all quartiles in **Figs 5E** and **5D**), in a proportion that was greater than among genes displaying chromatin-bound RNAs covering the exons (compare **Fig 5E to Fig 1B**). This suggested that the association between FluPol and RNA could not be solely attributed to the higher transcriptional activity of these genes.

Furthermore, gene ontology analysis on the genes at which PA was binding spliced chromatin-RNAs revealed an enrichment in host-cell defense pathways, consistent with the analysis of all upregulated genes (**Fig 5F**). Yet, we noted that the enrichment scores for Toll-like receptor (TLR) and IFN pathways were improved (**Fig 5F** and compare with **S1C Fig**). Finally, we noted an enrichment of the V5-PA subunit on the transcripts bound to the chromatin matched genes recruiting FluPol at their promoters, as documented by our DNA-ChIP assay, including the *OAS* genes, *DDX58/60*, *IFIT2/3/M1/M2*, and *ISG15/20* genes (**S5 Table**). Together, these observations suggested a degree of specificity in the targeting of FluPol, with a preference for genes upregulated by the infection, rather than genes highly expressed before and after the infection. In particular, we found that the IFNα/γ pathway was a target of FluPol, indicating that the binding of FluPol to nascent and possibly partially mature, mRNAs may play a role in the viral strategy to suppress the host-cell defense mechanisms.

### FluPol is associated with chromatin-bound RNAs originating from gene termination sites and downstream regions

We next used the V5-PA RNA ChIP-seq data to examine in more details the association of the FluPol with DoGs. The distribution of RNAs bound to the V5-PA subunit suggested an enrichment at the 3’end of genes (TES) and at regions covered by DoGs (**Fig 5C**). This was verified quantitatively by counting reads in the regions extending 40kb downstream of the TES of all genes, in bins of 10 kb (Left panel, **Fig 6A)**. When consolidating reads over the entirety of the 40kb-regions, the median density of V5-PA-associated RNAs was 2-fold higher than that observed in the negative control (right panel, **Fig 6A**). Within the set of V5-PA-associated DoGs, the number of upregulated genes was 3-to 4-fold higher than the number of unmodified DoGs (compare the brown and black bars in **Fig 6B**), while there were very few, if any, downregulated genes. Importantly, the proportion of increased DoGs in the V5-PA-bound RNA ChIP-seq data were greater than the proportion of increased DoGs in the chromatin RNA-seq data (compare **Fig 6B** with **Fig 2C**), and V5-PA-bound DoGs were greatly enriched at upregulated DoGs (47%-55%) compared to unmodified DoGs (6%-12%) or downregulated DoGs (0-7%) (**Fig 6B)**. This strongly suggested that the FluPol may participated in the viral complex involved in the inhibition termination processes and could be an active player in the mechanism causing the production of DoGs in infected cells.

**Figure 6:**
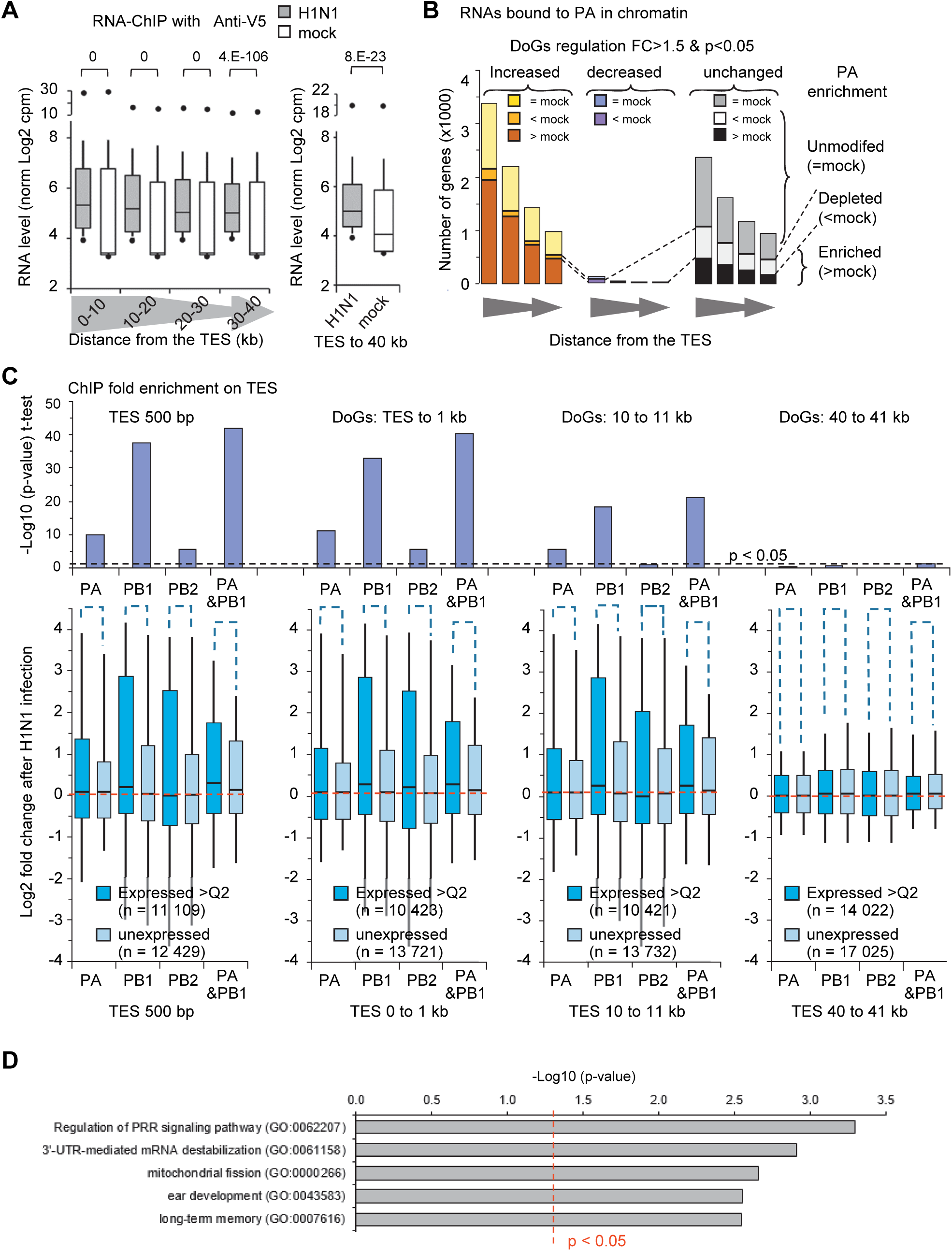
Enrichment of FluPol subunits on chromatin downstream of the 3’ ends of genes. A) RNA bound to the chromatin of infected A549 cells and immunoprecipitated with the indicated antibodies was analyzed by deep sequencing. For each selected gene, the indicated 10-kb windows from the TES were quantified for their DoGs levels evaluated in Log2(cpm). Data are shown in box plots as in **Fig 2B**. The right panel is the same quantification for all pooled windows..**B)** Counts of V5-PA enriched DoG regions and their differential transcription levels upon infection. Regions were considered enriched in PA (>mock, brown and black) when the average of the ChIP V5-PA signals minus the deviation was greater than twice the level in mock cells. Regions were considered depleted in PA (<mock, orange, purple, white) when the PA signal plus deviation was weaker than twice the signal obtained in mock cells. DoG levels were considered modified for a p-value <0.05 (paired t-test, two-tailed) and a fold change >1.5.. **C**) Fold change in enrichment for categories of the indicated terminal region relative to TES. These were the average differences between log2 cpm from infected and uninfected cells. Statistical analysis was performed on log2 fold change using Student t-test (two-tailed) to compare DoGs regions of unexpressed genes (light blue) and genes expressed above the median (blue). The dashed red line indicates no change. The p-values are shown above the box plots. The dashed line indicates the limit of p-value=0.05. In each case, the number (n) of TSS tested is indicated. PA&PB1 analyses were performed by combining their ChIP-seq signals. **D**) Pathway analysis of 1084 expressed genes (greater than the first quartile) for which the 1 kb windows downstream of the TES were significantly (p<0.05) enriched at least 2-fold for PA+PB1 between infected and control cells. P-values of Gene Ontology (GO) biological process 2021 were evaluated by Enrichr, and the threshold of p<0.05 is indicated by the dotted red line. The first five pathways are shown below.

To further document the presence of the FluPol in the DoG regions, we mined the DNA ChIP-seq data obtained with the antibodies against FluPol subunits, to quantify enrichment of individual subunits at TESs and at downstream regions. To avoid interference from overlapping, neighboring or converging genes, we selected genes separated from each other by at least 10 kb. PA and PB1 were most efficiently enriched at downstream regions of expressed genes, using silent genes to define the background signal (**Fig 6C**). The recruitment of these FluPol subunits was in average strongest at regions proximal to the TES (compare quantifications at 500bp, 1kb, 10kb and 40 kb away from the TES), suggesting that coding regions and proximal terminal downstream region were preferential sites for FluPol binding to nascent transcripts.

Gene ontology analysis on genes at which the FluPol was enriched on the 3’-ends revealed an enrichment in genes involved in cell defenses, in particular in the pathways of the pattern recognition receptors (with *BIRC2, GFI, ZDHHC5*) and mRNA stabilization (with *MOV10, DHX36*) (**Fig 6D, S6 Table).**

## Discussion

Examining the chromatin compartment has allowed us to gain clearer appreciation of the extensive impact of IAV infection on the host transcription machinery, altering both gene activity (**Fig. 1C**) and RNAPII distribution (**Fig. 2**). Monitoring of the distribution of the 3 subunits of FluPol on the chromatin of infected cells further suggested that the viral RNA polymerase could be a significant contributor to these alterations. This inhibition of gene termination results in a general decrease of the RNAPII at promoters and in gene bodies, and an increase in intergenic regions (**Figs 1D-F, S1D-S1G Figs**).

By using the presence of the 3 subunits of the viral RNA polymerase in the chromatin of infected cells, we showed that FluPol is not solely associated with promoters or enhancers where it is thought to snatch the caps of RNAPII-initiated RNAs (**Figs 3, 4)**. In fact, our observations suggest that FluPol remains associated with RNAPII during elongation, at transcriptional termination sites and also in downstream regions where RNAPII continues to run in intergenic regions (**Figs 5, 6**). FluPol was not only bound to the DNA of transcriptionally active genes but remains associated with host nascent transcripts (**Figs 5C-E, 6AB**), with a concentration to DoGs. To our knowledge, this is the first time that FluPol has been localized on a large scale across the genome and observed along the gene bodies and beyond. The association with DoGs suggests that FluPol could, in addition to NS1, also participate in the process of transcriptional termination inhibition.

Analysis of the regulatory pathways defined by the genes for which FluPol is most strongly enriched indicates that FluPol particularly targets genes of the anti-viral innate response. The participation of FluPol in the inhibition of transcriptional termination particularly on cellular defense genes would add a function of FluPol, in addition to cap-snatching, to participate in the viral escape strategy.

### FluPol genomic recruitment favors targets involved in cellular defense

Pioneering approaches to study the “cap snatching” process genome-wide have suggested that FluPol passively takes advantage of the most abundantly expressed genes [5]. Our observations suggest however that the targeting of FluPol may involve a more intricate process that includes some level of gene selectivity not solely dependent on expression levels. In this regard, a significant finding in our study is that FluPol is preferentially recruited to promoters undergoing H3K27 acetylation upon infection. This epigenetic mark serves as an indicator of increased promoter activity, implying that IAV exploits the host’s transcription activation machinery for FluPol recruitment. Interestingly, as this recruitment translates into transcriptional repression, the virus achieves a decoupling of the acetylation mark from gene activity. This serves as a remarkable demonstration of viral adaptation. Since the cell is naturally inclined to activate defense genes during an attack, selectively targeting these activated genes provides specificity towards defense-related genes. Consequently, FluPol is consistently found accumulating at TSSs (**Fig 3E, S2C Fig**), gene bodies (**Fig 5F**), TESs (**Fig 6D**) or enhancers **(Fig 4D**), associated with genes enriched in pathways related to the innate immune response.

This FluPol selectivity may be attained through interactions with host transcription factors induced by the infection, as for example IRF1 (**Fig 3E**). But FluPol may also target more general component of the transcription machinery, such as co-regulators previously identified by mass-spectrometry [16–19]. Further studies will be needed to fully understand this potential mechanism of FluPol recruitment.

### Transcriptional changes during viral infection confound the analysis of differential gene expression

It was known the IAV-induced terminal defects allowed the production of ‘‘downstream-of-gene’’ transcripts (DoGs) [11,30]. These DoGs, which extend beyond the poly-A termination signal, are typically transient RNA species and are usually not observed at high levels in standard RNA-seq experiments. However, in the case of IAV-infected cells, the DoGs and the associated pervasive transcription become significantly elevated, posing a challenge for accurate mRNA quantification using RNA-seq data. This challenge lies in the extensively modified distribution of the RNA-seq reads upon infection, a phenomenon known to partially defeat the assessment of differential gene expression by algorithms such as DeSeq2 [38,39]. In fact, the large redistribution of the RNAPII relocalization after the viral infection may also bias the evaluation of RNAPII presence in gene bodies as evaluated by ChIP-seq methods (**S1G fig** and [31]). We therefore confirmed by ChIP-PCR that accumulation of RNAPII is decreased within gene bodies upon IAV infection (**Fig 1E**). Our proposed RNA-seq normalization method, which involved utilizing reference genes with mRNAs possessing a half-life significantly longer than the duration of the infection, and hence assumed to maintain constant levels, unveiled an assessment uncertainty with DeSeq2 for approximately 25% of the genes designated as differentially expressed (**Fig 1C**), with many errors arising from the confusion between gene-specific reads and reads accounting for DoGs originating from upstream genes (**Figs 1D, 1F, S1D Fig**). This highlighted the utmost significance of selecting an appropriate normalization method when studying biological phenomena, such as viral infection by IAV, which not only impacts the expression level of genes, but also disrupts the distribution of RNAs (and RNAPII) across the genome. Lastly, it is worth noting that assessment of chromatin-bound RNAs allowed the observation of many transcriptional changes well-aligned with those obtained with more intricate techniques such as NET-Seq [11].

### FluPol’s chromatin association suggests Cap snatching is not limited to TSSs

Although we noted a certain inefficiency in ChIP with antibodies directed against the PA, PB1 subunits and even worse with the anti-PB2 antibody, the observed enrichment of FluPol on transcriptionally active genes versus silent genes documented the reliability of the data (**Fig 3, 6C**). In addition, the gene-wide distribution of FluPol was further supported by the RNA-ChIP using an anti-V5-tag antibody designed to recognize a V5-tagged variant of PA introduced into the virus (**Figs 5A 5B**).

Interactions between FluPol and the Ser5 phosphorylated form of RNAPII [8–10], as well as interaction with transcriptional coactivators [16–19] and the chromatin reader CHD1 that recognizes the promoter-specific H3K4me3 marks [20] have suggested that the process of cap-snatching was likely to occur when nascent RNAs exit from RNAPII paused at promoters [44]. While our examination of FluPol localization on chromatin did not challenge the recruitment of FluPol at the promoter level, it did cast uncertainty on the necessity of cap-snatching exclusively from polymerases awaiting promoter escape, and significantly expanded the spatial context in which this reaction could take place.

PA-associated RNAs are more often found up-regulated and significantly involved in the cellular response when they were exonic RNAs than when considering RNAs spanning the entire gene body (**Figs 5D 5E**). This suggested a potential association of PA with spliced RNAs in addition to the DoGs at the 3’-end of genes. Based on this and on the observations mentioned above, we propose a model where FluPol is recruited to TSSs but remains attached to RNAPII during elongation. As a result of co-transcriptional splicing, the pre-mRNA will undergo a least partial maturation by the time RNAPII and the accompanying FluPol reach the 3’ end of the transcribed gene. At the TES, termination inhibition causes transcription to continue beyond the poly-adenylation site, FluPol remaining associated with the elongating complex and the spliced mRNA (**Fig 7**).

**Figure 7.**
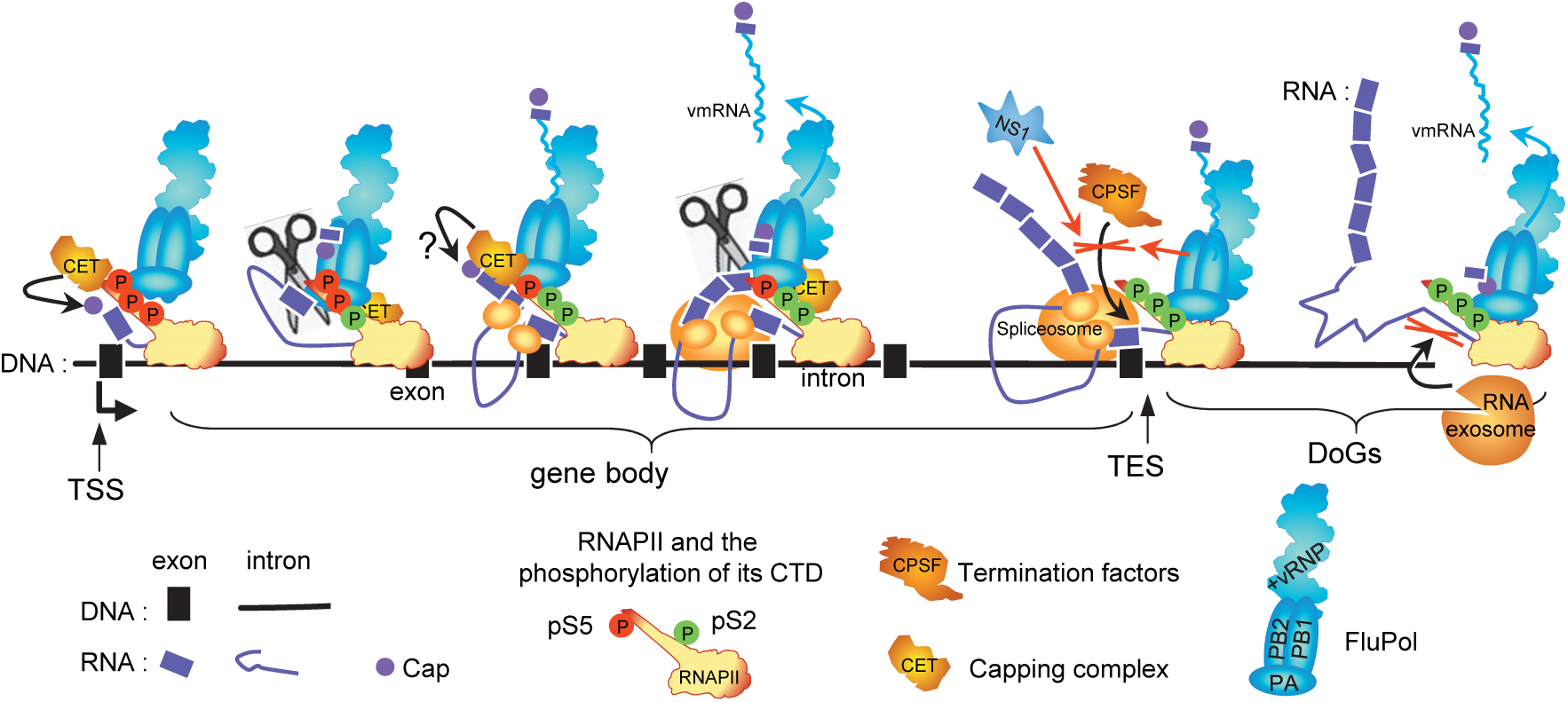
Working model of the functional association of FluPol with RNAPII. The FluPol is initially recruited at the level of the TSS and PB2 can bind the cap of the nascent RNA. During elongation, FluPol can remain bound to RNAPII either through its interaction with the S5p of the CTD, which remain present throughout the elongation process, through other RNAPII subunits (such as RPB4) or through the binding by the nascent transcript. Splicing of the nascent transcript occurs during elongation, producing intermediate spliced forms that remain associated to FluPol through the RNAPII in elongation. During these steps, potential re-capping through the capping enzyme and methyltransferase (CET) may provide new capped RNA primers for the FluPol. At the 3’ end of the genes, FluPol could be involved in the inhibition of termination complex (CPSF is also targeted by NS1) and also in the inhibition of the RNA exosome, thereby promoting the stabilisation of DoGs. The cap-snatching reaction of the nascent RNA by the PA subunit (scissors) could occur at any time during these steps. The nascent RNA remains bound to RNAPII in elongation until the 3’-end region, where RNAPII continues after TES due to termination inhibition. This proposed model could explain why FluPol is enriched at DoGs and exonic RNAs on chromatin.

This extended window of opportunity to steal caps may account for the preferential stealing of mRNA caps over those of enhancer RNAs or promoter antisense RNAs (PROMPT or antisense TSS) [22].

Indeed, as we have shown the presence of FluPol on enhancers (**Fig 4**), it was conceivable that enhancers, much more numerous than promoters could constitute an important source of capped transcripts to initiate vmRNA transcription. Apparently, this is not the case since eRNAs would represent only 3-5% of these sources compared to more than 70% for protein coding transcripts [22]. Similarly, it is surprising to note that although there are as many sites producing PROMPTs as sites producing mRNAs, i.e. promoters, PROMPTs are 12-to 40-fold less used as cap sources than mRNAs. Alternatively, the reduced stability of these eRNA/PROMPTs may reduce their availability for cap-snatching, although we note that inhibiting the RNA exosome to stabilize these transcripts resulted in only a modest 2-fold increase in PROMPT usage, while the usage of eRNA and antisense TSS remained unaffected [22].

This suggests that transcript lifetime is not the major cause of mRNA-use as a source of cap by FluPol. This and our results regarding the presence of FluPol along the entire length of transcribed genes and even beyond, leads us to propose the hypothesis that if cap stealing occurs predominantly on mRNAs it is because it does not occur immediately after transcriptional initiation when RNAPII is paused at the promoter (otherwise the proportion of cap source between antisense and sense transcripts would be equivalent). The ability of FluPol to remain attached to the elongating RNAPII gives it the opportunity to act on the cap after transcriptional initiation, at the time, when steric hindrance between the transcriptional complex and the nascent RNA could be reduced. Such cap-snatching taking place at any stage of pre-mRNA transcription would align with the observation that cap-snatching occurs co-transcriptionally [22].

It is noteworthy that capping enzymes maintain their association with the RNAPII CTD during transcript elongation and even beyond the site of polyadenylation [24]. Additionally, the capping activity is not confined to promoters [45]. Thus, by remaining associated with the elongating RNAPII, FluPol may be offered multiple opportunities for cap-snatching since nascent transcripts can be theoretically recapped during elongation [46].

The interaction with pS5-RNAPII would then primarily facilitate FluPol’s entry onto the genes. It is also noteworthy that pS5-RNAPII does not disappear during elongation since this phosphorylation correlates with cotranscriptional splicing process [47]. In addition, other interactions with the transcription machinery, including RBP4 [48] would be required for the prolonged residence of FluPol on the transcribing gene and the improved probability of cap-snatching.

### FluPol’s association with the elongating RNAPII may help the virus evade cellular defense

A FluPol-RNAPII association spanning from transcription start to termination may provide the virus with multiple advantages. Firstly, it may improve access to host mRNA maturation end export factors, possibly facilitating the splicing of genomic segments 1, 7 and 8 (which are partially spliced to generate PB2-S1, M2, M42, NS3 and NS2 viral proteins), for terminal cleavage and polyadenylation [49] and for nuclear export [1,50]. Next, it may allow concentrating the virus-induced transcription termination defect on the genes involved in cellular defense. Indeed, the presence of FluPol on termination and downstream regions (**Fig. 5F, 6C**) and its association with DoGs (**Fig. 6A**) strongly suggest that FluPol participates in the inhibition of termination. Thus, a sustained association of FluPol with RNAPII may allow combining the selective recruitment of FluPol with termination defects, thereby impeding the expression of genes activated by the viral attack. We note also that an implication of both FluPol and NS1 in impairing RNAPII termination may explain why termination defects that are still observed in IAV mutants lacking NS1 expression [11].

Reducing the pool of RNAPIIs available for transcription re-initiate at defense genes may, however, not be the sole benefit of DoGs. Because the intergenic regions crossed by DoGs are rich in repeated sequences (SINE, LINE, LTR), transcription beyond gene boundaries will result in the production of RNA species prone to fold into double-stranded structures and potentially recognized by intracellular PRRs [51]. The robust increase in DoG accumulation during viral infection, along with the transcriptional repression of genes related to the PRR pathway, may therefore result in the saturation of the double-stranded RNA detection system. In that context, we have recently shown DOG accumulation is also observed in association with cellular senescence, along with accumulation of promoter RNAs [52]. In this case, this accumulation of intergenic transcripts appeared to be a consequence of their reduced degradation rather than their increased production, with a clearly reduced expression of one or more subunits of the RNA exosome. This latter complex, involved in the degradation of a large range of RNA species, has been reported to associate with FluPol for the benefit of viral multiplication [22]. This positive effect may in part be mediated by an inhibitory effect of FluPol on the activity of RNA exosome. Collectively, these different processes may facilitate the establishment of an optimal environment for viral activities, including the formation of cellular:viral RNA hybrids that are indispensable for unhindered viral transcription.

## Material and Methods

### H1N1 virus construction and preparation

This study uses Influenza virus A/WSN/1933 (H1N1). Wild-type and V5-tagged PA versions of the viruses were generated by reverse genetics using the 12-plasmid reverse genetics system as described previously [53]. The V5-epitope (amino acid sequence NH2-GKPIPNPLLGLDST) nucleotide sequence was fused in frame at the 3’ end of the PA gene through a three glycine-long linker. To generate the PA-V5 plasmid used for virus rescue, a silent Nco1 restriction site was first introduced at nucleotide position 2116-2121 to allow introduction of exogenous sequences in this PA 3’ domain. To preserve the PA viral incorporation signal, the PA-V5 encoding sequence was followed by a stop codon and the last 116 3’-nucleotides of the PA segment.

### Cell culture, H1N1 infection

A549 cells (CCL-185) were purchased from ATCC. Cells were maintained in Dulbecco’s modified Eagle’s medium (Gibco) supplemented with 7% (v/v) fetal bovine serum (Thermo Scientific) and 100 U/ml penicillin-streptomycin (Gibco). A549 pellet of cells, infected (MOI=5, h.p.i=6h) with V5-tag-PA (virusV5) or normal virus WSN H1N1. Cells have been not fixed or fixed by 0.5%, 1%, 2% formaldehyde in PBS solution during 10 min at room temperature prior to harvest. Cells were pelleted and frozen at −80°C prior performing the extracts.

### Antibodies

Antibodies were purchased from Diagenode for H3K27ac (C15410196); H3K4me1 (C15410037) from Abcam for RNAPII (8WG16), RNAPII pS5 (ab5131) for Western blot, RNAPII pS5 (CTD 4H8) for ChIP, histone H3 (Ab1791), from ThermoFisher for PA (PA5-31315), PB1 (PA5-34914) and PB2 (PA5-32220); from Bethyl for anti-V5 epitope (A190-120A).

### Chromatin immunoprecipitation (ChIP and RNA-ChIP) and RNA-chromatin extraction

ChIP assays were performed as previously described [54,55]. Cells were crosslinked during 10 min at room temperature in PBS with 1% (v/v) formaldehyde, followed by a 5 min wash in PBS containing 125 mM Glycine before harvest. Resuspended cells (10-20E6 in 1.5 mL µtube) were incubated on ice 5 min in 1 mL of chilled buffer A (0.25% TRITON, 10 mM TRIS pH8, 10 mM EDTA, 0.5 mM EGTA), 30 min in 1 mL of buffer B (250 mM NaCl, 50 mM TRIS pH8, 1 mM EDTA, 0.5 mM EGTA), and then resuspended in 100 to 200 µL of buffer C (1% SDS, 10 mM TRIS pH8, 1 mM EDTA, 0.5 mM EGTA) at room temperature. All buffers were extemporately supplemented by 0.5 U/µL RNAsin (Promega), Complete protease and PhosSTOP phosphatase inhibitors (Roche). Cell suspensions in 0.6 mL µtube were sonicated in water bath at 4°C during 12 min (15 sec. ON, 15 sec. OFF) with a Bioruptor apparatus (Diagenode) setted on high power and then clarified by 10 min centrifigation at 10 000 rpm, 4°C. Shearing of the DNA was checked after reversing the crosslinking on agarose gel electrophoresis to be around 300-500 bp. Sheared chromatin was quantified by optical density (260 nm) and diluted 10-fold in IP buffer to final concentrations: 1% TRITON, 0.1% NaDeoxycholate, 0.1% SDS, 150 mM NaCl, 10 mM TRIS pH8, 1 mM EDTA, 0.5 mM EGTA, 1 U/µL RNasin, 1X protease and pshotase inhibitors). For ChIP-PCR assays 30 µg of chromatin and 2 µg of antibodies, in a final volume of 500 µL, were incubated at 4°C for 16 h on a wheel. For ChIP-seq or RNA-ChIP-seq assays, 200 µg of chromatin was incubated with 10 µg of antibodies in a final volume of 2 mL. 20 µL of saturated magnetic beads coupled to protein G (Dynabeads) per µg of antibody were used to recover the immuno-complexes. After 2 h of incubation the bound complexes were washed extensively 10 min at room temperature on a wheel in the following wash buffers: WBI (1% TRITON, 0.1% NaDOC, 150 mM NaCl, 10 mM TRIS pH8), WBII (1% TRITON, 1% NaDOC, 150 mM KCl, 10 mM TRIS pH8), WBIII (0.5% TRITON, 0.1% NaDOC, 250 mM NaCl, 10 mM TRIS pH8), WBIV (0.5% Igepal CA630 (Sigma), 0.5% NaDOC, 250 mM LiCl, 10 mM TRIS pH8, 1 mM EDTA), WBV (0.1% Igepal, 150 mM NaCl, 20 mM TRIS pH8, 1 mM EDTA), WBVI (0.001% Igepal, 10 mM TRIS pH8). 20 µg of sheared chromatin used as input and ChIP beads were then boiled 10 min in 100 µL H2O containing 10% (V/W) chelex resin (BioRad), followed by Proteinase K (0.2 mg/mL)-digestion for 30 min at 55°C, and then finally incubated 10 min at 100°C. 0.5 µL was used for quantitative real-time PCR assays. For RNA-chromatin extraction, supernatant from the sonicated step were purified by acidic phenol/chloroform method and ethanol-precipitation with 5 µg of Glycoblue (Ambion) as carrier. After resuspension these input, as well as the eluate from RNA-ChIP were treated by 5 U of TURBO-DNase (Ambion) 1 h a 37°C prior purification by acidic phenol/chloroform method and ethanol-precipitation with 5 µg of Glycoblue (Ambion) as carrier.

These samples were then checked for the absence of DNA contamination by qPCR with 1 µL of eluate and primers targeting genomic repetitive elements. Another round of DNAse digestion and purification was performed in case of qPCR signal < 35 Ct.

### Library preparation and high throughput sequencing

1 to 5 nanograms of immunoprecipitated DNA were used to build DNA libraries. Libraries were prepared following the instruction from NEBNext® Ultra™ II DNA Library Prep Kit for Illumina® (NEB #E7645L) and indexes (NEB #7335S; #E7500S; #E7710S). No sized selections were applied. The quality and size of the libraries were assessed using the 2100 Bioanalyzer (Agilent). The amount of the DNA used for library preparation and the amount of DNA libraries were measured using Qubit (Invitrogen). Indexed libraries were pooled according to the Illumina calculator (https://support.illumina.com/help/pooling-calculator/pooling-calculator.htm) and deep sequenced by the Illumina platform provided by Novogene (HiSeq-SE50). RNA from V5 ChIP in non-infected (mock) cells in triplicate were pooled to ensure sufficient material equivalent to individual V5-PA ChIP in infected cells. Libraries from ChIPed RNA and input chromatin-RNA were performed by the GenomEast platform, a member of the ‘France Genomique’ consortium (ANR-10-INBS-0009) (IGBMC, Illkirch, France). RNA-Seq libraries were generated from 100 to 300 ng of RNA using TruSeq Stranded Total RNA Library Prep Gold kit and TruSeq RNA Single Indexes kits A and B (Illumina, San Diego, CA), according to manufacturer’s instructions. Briefly, ribosomal RNA was removed by Ribo-Zero beads. After chemical fragmentation depleted RNA were reverse-transcribed with random primers. After second strand synthesis by DNA Polymerase I and RNase H and blunting, the cDNA were ligated to adapters and amplified by PCR for 12 cycles. Residual primers were removed by AMPure XP beads and the libraries were sequenced on Illumina Hiseq 4000 sequencer as Paired-End 100 base reads. Image analysis and base calling were performed using RTA 2.7.7 and bcl2fastq 2.17.1.14.

### Data availability

ChIP-seq, RNA-ChIP-seq, RNA-seq data have been deposited in the NCBI Gene Expression Omnibus database under GEO accession number GSE218084: (access for reviewer: token ojmbsogibtgrlyl into the box).

### Protein extraction for immunoprecipitation

Pellet of frozen cells corresponding to 10E6 cells were thawed in ice and resuspended in 500 µL of Buffer A with protease inhibitors (Roche). 150 µL of suspension were supplemented with 180 µL of buffer B and sonicated 5 min at 4°C by BioRuptor (high power, 15 sec. ON / 15 sec. OFF). Clarified supernatants were quantified by DO (260 nm). 20 µg of protein extract were incubated with 1 µg of indicated antibodies during 16 h at 4°C on a wheel and then 2 h with 30 µL of anti-rabbit Dynabeads. Beads were washed 5 times in wash buffer WBV, resuspended in laemmli buffer with 100 mM DTT and boiled 20 min and supernatant was analyzed by western blot.

### Western blot

Denaturated proteins were resolved by SDS–PAGE (4–12% Criterion XT Bis-Tris Protein Gel; Bio-Rad), and transferred to nitrocellulose membrane (Bio-Rad). Blocked membrane were incubated with 1/1000 diluted first antibody as indicated in figures in PBS with 0.1% Tween-20 and 5% (V/W) non-fat milk, washed 10 min 3 times in PBS-tween(0.1%), incubated 1 h at room temperature with 1/3000 anti-Mouse StarBright blue700 (BioRad) or 1/2000True blot HRP anti-rabbit antibodies. After 4×10 min washes in PBS-tween(0.1%), the membranes were revealed revealed by chemiluminescence (for HRP), and quantified using Chemidoc MP imaging system (Bio-Rad).

### Real-time quantitative PCR

1 µL of ChIP eluate was used for quantitative real-time PCR (qPCR) in 10 μL reactions with Brillant III Ultra Fast SYBR-Green Mix (Agilent) using a Stratagene MX3005p system. The analysis of qPCR was performed using the MxPro software. The sequences of primers used for PCR are in **S7 Table**. Statiscal analysis and graphs were produced with Microsoft Excel.

### Bioinformatic analysis

The human genome of reference considered for this study was hg19 homo sapiens primary assembly from Ensembl. After reads alignment, the SAM files were then converted to BAM files and sorted by coordinate and indexed using samtools (v1.7) [56]. For ChIP-seq data from this study, from those in MDM cells (GSE103477) (Heinz et al. 2018), or from A549 H3K4me3 ENCODE track (GSE91218), reads were mapped to hg19 using bowtie2 (v2.3.4) [57] (parameters: -N 0 -k 1 –very-sensitive-local). We then selected reads with a MAPQ equal or higher than 30 corresponding to uniquely mapped reads for further analysis. After indexing, MACS2 (v.2.1.1) [58] was used to call peaks (parameters: -p 0.05). For RNA-chromatin and RNA-ChIP-seq data, mapping was carried out with STAR (v2.6.0b) [59] (parameters:--outFilterMismatchNmax 1--outSAMmultNmax 1--outMultimapperOrder Random--outFilterMultimapNmax 30). Data observations were performed with the Integrative Genomics Viewer software [60] using the CPM normalized BigWig files produced from the BAM files by Deeptools suite (parameter:--normalizeUsing CPM) [61]

Evaluation of gene expression from chromatin-RNA (our study) was performed by using featureCounts (v1.28.1) [62] from the Rsubreads package (parameter:-s 2). These read counts were analysed using the DESeq2 [63] package in order to test for the differential gene expression (DGE). The normalization, the dispersion estimation and the statistical analysis were performed with DESeq2 using the default parameters. Raw *P*-values were adjusted for multiple testing according to the Benjamini and Hochberg (BH) procedure and genes with an adjusted *P*-value lower than 0.05 and a fold change greater than 50% were considered differentially expressed. The raw count from featureCount were also used to produce normalized gene levels using reference genes as follow: counts covering mRNAs were converted in rpkm by considering the library total count of each sample and the length of each gene as defined in Ensembl. The average level of genes with half-life greater than 15 h as defined in [37] and that were more expressed than 75% of all expressed genes (corresponding 417 genes with level higher than 300 rpkm for chromatin-RNA) was used to calculate a corrective index applied on the gene levels in infected cells. The maximun mRNA level between mock and infected cells were used to draw the MA-plot (**S1B Fig**). Genes with a *P*-value from a paired t test (n=3) lower than 0.05 and a fold change greater than 50% were considered differentially expressed.

Signal evaluation of ChIP-seq, RNA-ChIPseq and RNA-seq on defined windows were performed as followed. Multiples regions based on the Ensembl annotations (hg19, version 87) were generated using the bedtools suite (v2.27.1) [64], 500 bp around the center of the intron that is closest to the center of the gene, 500 bp around the TSS, 500 bp around the TES, from TES to 1 kb downstream gene, from TES to 10 kb downstream gene and 40 kb to 41 kb downstream gene. In the case of the downstream region of genes, only regions retained were those not overlapping and distant by at least 10 kb with with any other gene from the TES to the end of the counting region. For TSS regions, windows retained were those distant by at least 10 kb from both extremities of other genes.

Consecutive 10 kb windows were generated after TES of genes with a maximum of 40 kb as long as there was no overlap with downstream genes oriented in the same direction. All previously generated counting tracks were converted to the SAF format and then used as annotation file in featureCounts (v1.6.1) (parameter:-s 2 for RNA-ChIP-seq or -s 0 for ChIP-seq). Regions were considered differentially enriched between infected and mock cells as indicated in each figure.

To assess H3K27ac signal at enhancers potential locations in ChIP-seq of infected cells, we have firstly excluded H3K27ac peaks from 10 kb windows around promoters regions defined by A549 H3K4me3 peaks from ENCODE (GSE91218) by using bedtools intersect (v2.27.1). All retained H3K27ac peaks and successive regions of 1kb around the retained H3K27ac peaks were designed (up to 4kb maximum) and used as counting tracks for featureCounts (v1.6.1) (default parameters for paired-end data). Statistical analysis and graphs were produced with Microsoft Excel.

Pathway analysis with gene name were performed with Enrichr [65]. To assess Pathway analysis of genes near enhancers, the proximal five genes in the regions extending 500 kbp downstream and the proximal five genes in the regions extending 500 kbp upstream from the H3K27ac peaks were selected as described in the text. The H3K27ac content of loci were evaluated by normalizing the H3K27ac ChIP signal to the H3 ChIP signal. Genes without any reads covering the exons were not considered. Gene lists were then submitted to the Enrichr website (https://maayanlab.cloud/Enrichr/).

## Acknowledgments

We are grateful to our colleagues C. Rachez for helpful discussions, Catherine Bodin for technical assistance and Edith Ollivier, Aurelie Prats, Johanne deMarchi for administrative assistance and Jeanne Le Peillet (Beink) for graphic advises.

## Author Contributions

Conceptualization:.EB, CM

Funding acquisition:.BD, CM, EB

Investigation:. EB, JY, MC, NL, JM

Project administration: EB

Writing – original draft: EB

---

**Supplementary figure S1 (related to Fig 1):**
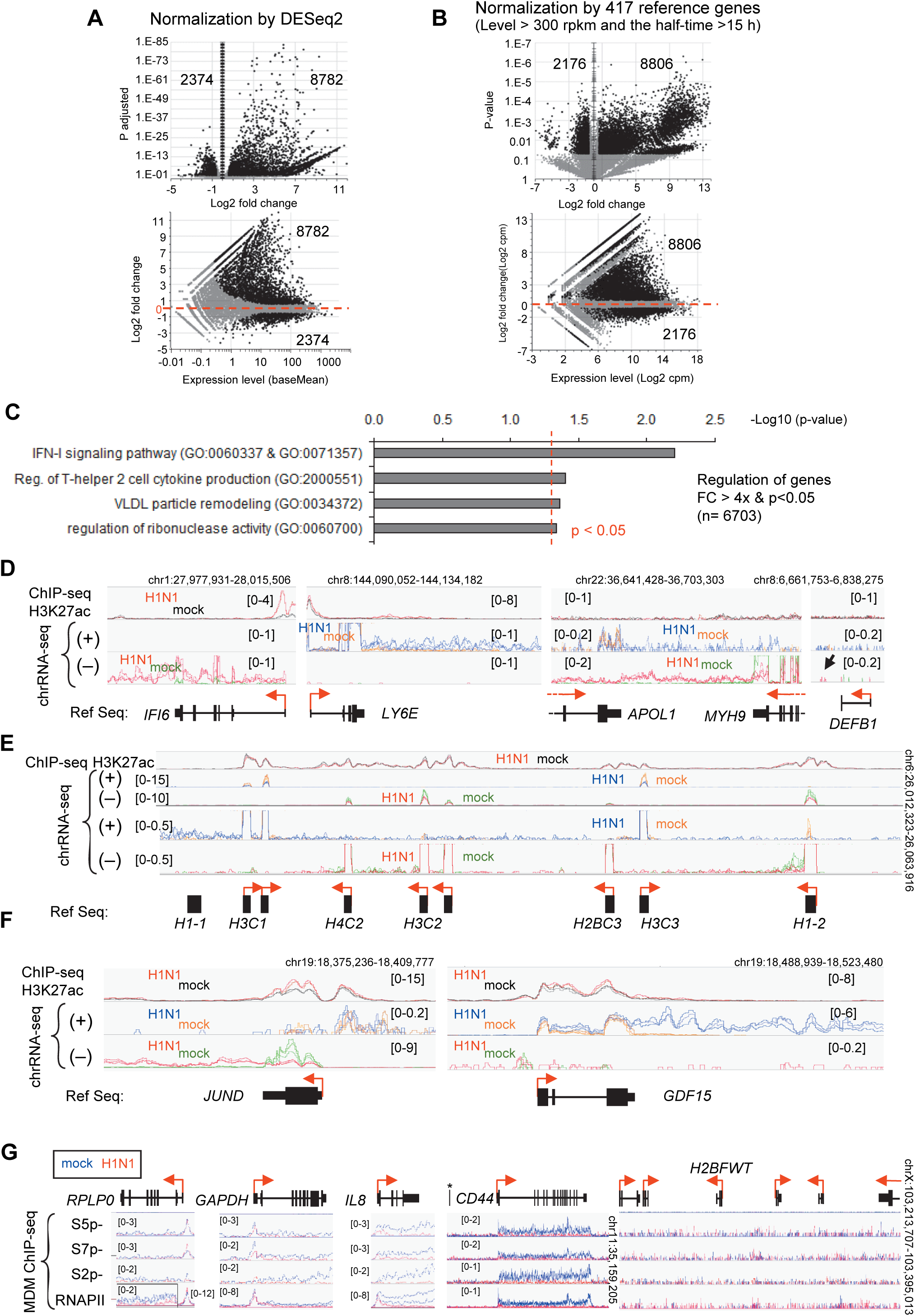
Differential gene expression at 6 h in H1N1-infected A549 cells versus uninfected cells. **A**) Differential RNA levels between H1N1 infected and uninfected cells using DESeq2 package. Variations were considered significant (black dot) if the mean fold change (FC) was >1.5 and the adjusted p-value was <0.05. The number of black dots is indicated. **B)** Differential levels of chromatin-associated RNA using normalization based on the average variation of 417 reference genes expressing higher levels (> 10 log2 cpm) of long half-life (> 15 h) transcripts [35].. Variations are considered significant (black dot) if the mean fold change is >1.5 and the p-value <0.05 calculated by paired t-test (two-tailed) on log2(cpm). **C)** Pathway analysis of 6703 genes upregulated 4-fold with p<0.05 between infected and mock cells. P-values of Gene Ontology (GO) 2021 were evaluated by Enrichr, and the threshold of p<0.05 is indicated by the dotted red line. **D, E, F**) Examples illustrating the variation in read distribution in H1N1 infected cells from IGV visualization of indicated loci. The RNA-seq independent triplicates have been overlaid in the same color as indicated for each strand orientation. The top trace shows the enrichment of H3K27ac on the chromatin of infected or mock cells. The scale range of the tracks is indicated in parentheses. The bottom track (Ref Seq) shows the position of the genes and their orientation as indicated by the red arrows. **G**) Distribution of indicated phosphorylated (at serine 5, 7 and 2) and total RNAPII on analyzed genes Fig 1E. ChIP-seq assays using antibodies against the indicated RNAPII were performed in infected (red line) or uninfected MDM cells (blue line) from the GSE103477 dataset [31]. The range scale are indicated in brackets.

**Supplementary figure S2 (Related to Fig 3):**
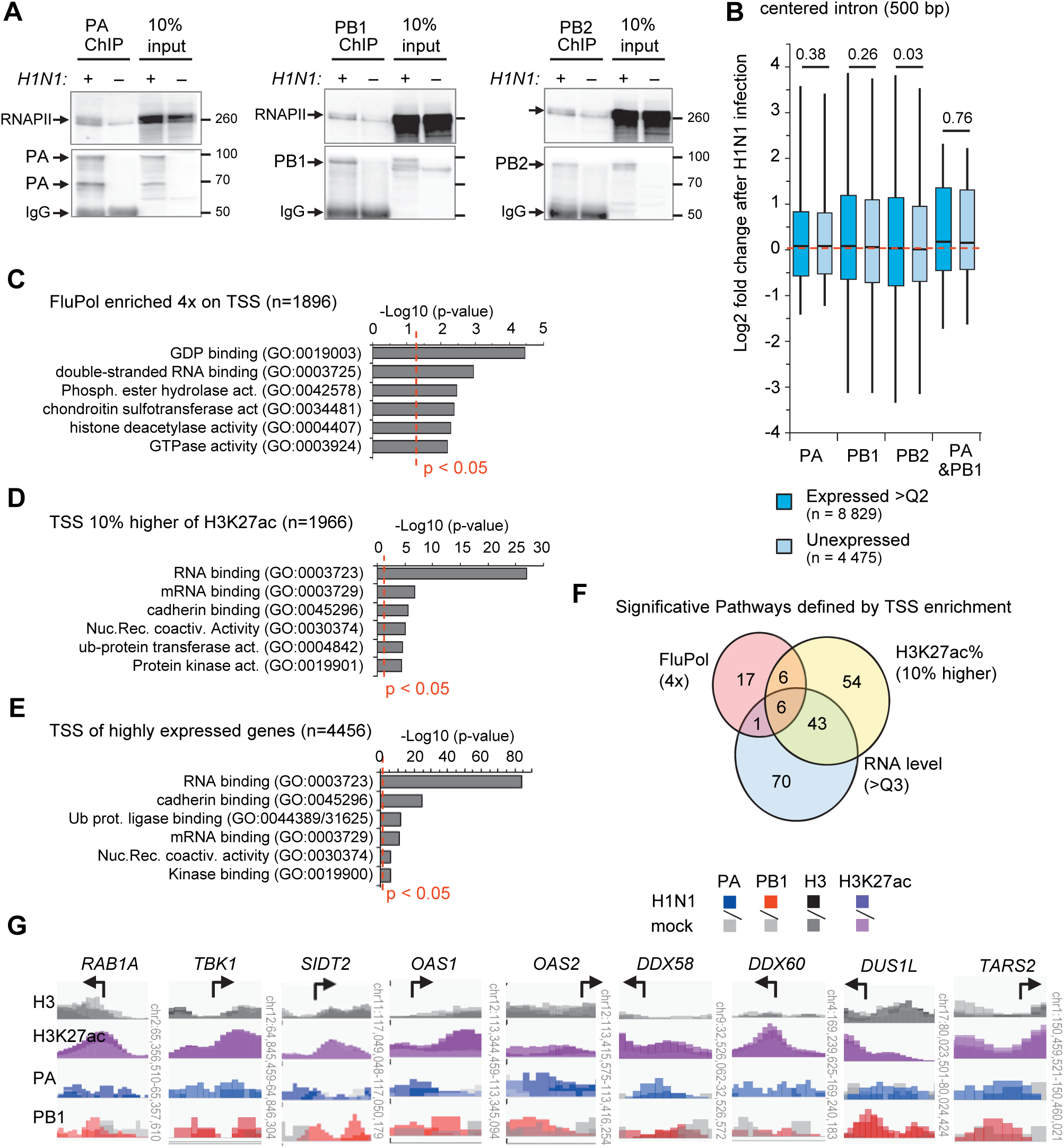
Immunoprecipitation efficiencies of antibodies directed against the Flu Pol subunits. **A**) Infected (+) or uninfected (-) cells were extracted and subjected to immunoprecipitation with the indicated bodies used in the ChIP assay. The eluates and 10% of the extracts (input) were resolved by Western blot. The lower part of the membranes (50 to 120 kDa) was detected with the corresponding antibodies and the upper part (120 kDa-top) with RNAPII antibodies. **B**) The same analysis, as in **Fig 3B**, was performed for the averages of intronic coverage. For each gene, the more central intron was used to define a centered 500 bp counting windows. Only windows separated by 10 kb from the TSS were considered. **C**) Pathway analysis of 1896 expressed genes (above the first quartile) for which the average ChIP-seq signal A or PB1 was enriched 4-fold on TSS (with H3K27ac/H3 levels above the first quartile) in infected vs. mock cells. Only TSS rated by 5 kbp from other genes were considered. P-values of Gene Ontology (GO) molecular function 2021 were evaluated by Enrichr, and the threshold of p<0.05 is indicated by the dotted red line. The first six pathways are shown and the full lists are in **S3A Table**. **D, E**) The same Gene Ontology analysis was performed on either the 1966 genes with 10% higher 27ac on their TSS or the 4456 genes in the upper quartile of RNA levels. **F**) Venn diagram of significant GO terms for molecular function (p-value <0.05) obtained in **S2C,D,E Figs**. **G**) Examples of 500 bp regions around the TSS of the indicated genes selected in the pathways in **C**). The ChIP-seq independent replicates performed with antibodies against the indicated proteins in infected or mock cells are overlaid in the indicated color. The black arrows indicate the position of the TSS and the orientation of the gene.

**Supplementary figure S3 (Related to Fig 5):**
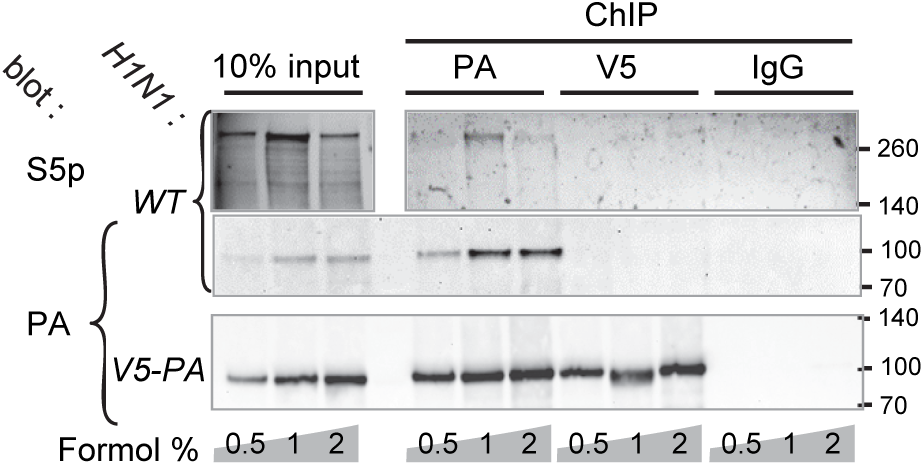
Immunoprecipitation efficiencies of Antibodies directed against the Flu Pol subunits. A549 cells infected cells by WT H1N1 or V5-PA engineered H1N1 were crosslinked by indicated % of formaldehyde or not. Chromatin fractions were subjected to immunoprecipitation with PA, V5 antibodies or non-immune IgG. Eluates and 10% of extracts were resolved by Western blot. The lower part of the membranes was visualized with PA antibodies and the upper part with phospho-RNAPII antibodies.

## Supplementary Tables 1-7

Excel spreadsheet containing, in separate sheets, the underlying numerical data, statistical analysis and Lists of genes used in pathway analysis for Figs

**TabS1**: RNAs bound to chromatin and differential RNA level (related to **Figs 1A-C, S1A and S1B**) **TabS2**: Gene Ontology Analysis of 6703 genes upregulated >4 (p<0.05) (related to **S1C Fig**) **TabS3A**: Gene lists and Gene Ontology pathway for Molecular function for 1896 genes on which PA or PB1 were enriched at least 4 fold on TSS, for 1996 genes with the highest level of H3K27ac on TSS, and for 4456 genes with high RNA levels (related to **S3C-F Figs**)

**TabS3B**: Enrichment of FluPol target genes, of H3K27ac-upregulated genes, and of genes with RNA level change with consensus of indicated transcription factors in ENCODE and ChEA (related to **Fig 3E**).

**TabS4A**: Pathway Analysis of 1966 expressed genes close to enhancer with more 20% FluPol enrichment (related to **Fig 4D**)

**TabS4B**: Pathway Analysis of 3208 expressed genes close to enhancer without FluPol enrichment (related to **Fig 4D**)

**TabS4C**: Pathway Analysis of 603 expressed genes close to enhancer with higher enrichment of FluPol (related to **Fig 4D**)

**TabS5**: Pathway analysis of 1696 genes where v5-PA bound RNAs were enriched at least 4-fold (related to **Fig 5F**)

**TabS6**: Pathway analysis of 1084 genes on which PA and PB1 were enriched at least 2 fold on TSE (related to **Fig 6D**)

**TabS7**: list and sequences of qPCR primers

